# Post-transcriptional modulation of Dscam1 enhances axonal growth in development and after injury

**DOI:** 10.1101/148239

**Authors:** Marta Koch, Maya Nicolas, Marlen Zschaetzsch, Natalie de Geest, Annelies Claeys, Jiekun Yan, Matthew Morgan, Marie-Luise Erfurth, Matthew Holt, Dietmar Schmucker, Bassem A Hassan

## Abstract

Injury to the adult central nervous systems (CNS) results in severe long-term disability because damaged CNS connections rarely regenerate. Although several axon regeneration regulators have been proposed, intrinsic regenerative mechanisms remain largely unexplored. Here, we use a *Drosophila* CNS injury model to identify a novel pro-regeneration signaling pathway. We conducted a genetic screen of approximately three hundred candidate genes and identified three strong inducers of axonal growth and regeneration: the Down Syndrome Cell Adhesion Molecule (Dscam1), the de-ubiquitinating enzyme Fat Facets (Faf)/Usp9x and the Jun N-Terminal Kinase (JNK) pathway transcription factor Kayak (Kay)/Fos. Genetic and biochemical analyses link these genes in a common signaling pathway whereby Faf stabilizes Dscam1 protein levels, by acting on the 3’-UTR of its mRNA, and Dscam1 acts upstream of the growth-promoting JNK signal. The mammalian homolog of Faf, Usp9x/FAM, shares both the regenerative and Dscam1 stabilizing activities, suggesting a conserved mechanism.

## Introduction

During CNS development axons grow in a tightly regulated manner to generate an intricate and complex pattern of neuronal connectivity. In most animal species, injury to the adult CNS, either by physical trauma or in the context of neurodegeneration, has devastating long-term consequences-in part because of the inability of mature neurons to regenerate severed axons, in contrast to peripheral nervous system (PNS) neurons. Functional regeneration requires damaged axons to first start re-growing and then to continue to navigate through a strongly inhibitory environment, before they can reach their synaptic partners and establish functional connections. Therefore, both the presence of extrinsic inhibitory factors as well as a lack of intrinsic growth capacity prevent axonal regrowth in the injured CNS (Kaplan et al., 2015). Targeting extrinsic inhibitory factors has so far led to limited regeneration of injured axons (Cafferty et al., 2010; Lee et al., 2010), suggesting that creating a permissive environment is not sufficient to allow regeneration. Therefore, elucidation of the molecular mechanisms underlying the early steps of intrinsic axonal regrowth after CNS damage is critical to fully understand the regeneration process. Even though neural circuits retain a remarkable degree of synaptic plasticity in adulthood, the mature CNS can no longer support the robust axonal growth that was once required to establish neuronal connectivity during development, suggesting that the neuronal intrinsic growth ability is largely lost. Indeed, mammalian CNS axons show a higher regenerative capacity during earlier stages of development, illustrating the importance of intrinsic factors to CNS regenerative failure (Liu et al., 2011; Shimizu et al., 1990). In PNS neurons, axonal injury results in a regeneration program that shares key molecular features with developmental axon growth (Harel and Strittmatter, 2006; Makwana and Raivich, 2005; Raivich and Makwana, 2007; Yaniv et al., 2012). In particular, the JNK pathway has emerged as a conserved signal for axonal growth and regeneration in the CNS and PNS in mammals, flies and worms (Arthur-Farraj et al., 2012; Ayaz et al., 2008; Li et al., 2012; Nix et al., 2011; Raivich et al., 2004; Raivich and Makwana, 2007). This suggests that conserved developmental axonal growth signaling pathways may be key targets to boost efficient regeneration after injury.

Studies in mice have made unique contributions to our understanding of the molecular basis of axonal regeneration. Nevertheless, the experiments are costly and time-consuming and necessitate a gene-by-gene approach. More recently, simpler genetic model organisms such *C. elegans* and *Drosophila* have proven useful to identify and study novel genes involved in axonal regrowth after injury (Ayaz et al., 2008; Chen et al., 2011; Fang and Bonini, 2012; Fang et al., 2012; Gabel et al., 2008; Kato et al., 2011; Leyssen et al., 2005; Yanik et al., 2004). Interestingly, unlike *C. elegans* neurons and developing *Drosophila* neurons, injured adult *Drosophila* CNS axons fail to regrow after injury, much like their mammalian counterparts (Ayaz et al., 2008). Furthermore, adult *Drosophila* CNS axons show remarkable morphological and genetic hallmarks of mammalian axonal responses to injury, including the formation of retraction bulbs, Wallerian degeneration of the distal fragment, transient upregulation of JNK, and regeneration upon activation of protein kinase A and JNK signaling (Ayaz et al., 2008; Leyssen et al., 2005; MacDonald et al., 2006). This makes the *Drosophila* adult CNS a particularly powerful model system to systematically search for novel axonal regeneration genes.

Here, we perform a two-step genetic screen of ~300 genes selected by GO term, and identify 13 that promote axonal outgrowth during development in post-mitotic CNS neurons. We then test those genes in an adult *Drosophila* model of CNS injury. Using this approach we identify three robust axonal regeneration regulators, which we find to interact in a novel axonal growth and regeneration signaling pathway. Specifically, the deubiquitinating enzyme Fat facets (Faf) promotes axonal regrowth after injury via the Down syndrome cell adhesion molecule (Dscam1). Faf stabilizes Dscam1 by acting on Dscam1 3’-UTR through the kinase DLK1/Wallenda (Wnd). Faf and Dscam1 act upstream of JNK signaling and its nuclear effector Kayak (Fos). The functional role of Faf in promoting axonal regeneration appears to be conserved in mammals as demonstrated by the ability of the mouse homologue of Faf, Usp9X/FAM to also stabilize Dscam1 and promote axonal regrowth in the injured fly CNS.

## RESULTS

### A genetic screen for axonal growth in development and after injury

To perform a screen for axonal growth and regeneration (Fig. 1a,b), we selected genes which: 1) are associated with the Gene Ontology (GO) terms neural development and neurite morphogenesis, 2) have Gal4 inducible transgenes available and 3) represent a diversity of molecular functions, including receptors, protein turnover, transcription factors, and chromatin modifiers (Supplementary data file 1). 307 genes matching these criteria were first tested for their ability to induce developmental axonal over-growth in small Lateral Neurons ventral (sLNv), a small cluster of neurons with a highly stereotyped axonal morphology which can be readily quantified (Helfrich-Forster et al., 2007; Leyssen et al., 2005) and that has been previously used to investigate the molecular mechanisms underlying regeneration in the fly CNS ^12^ (Fig. 1 a, Fig.2 a and b). The post-mitotic *Pdf-Gal4* driver was used to express GFP together with each of the selected genes, and the length of axonal growth was quantified in comparison with controls. Expression of 13 genes (4.2%) promoted significantly increased axonal growth with no obvious adverse effects on neuronal survival or axonal trajectory (Fig. 2 a-e). In a second selection step (Fig. 1 b), these 13 genes were evaluated in an acute sLNv axonal injury model in *Drosophila* brains explanted and kept in culture (Ayaz et al., 2008; Koch, 2012). Given their superficial location, sLNv axons are easily accessible for injury, and were physically severed using an ultrasonic microchisel. Using the temperature dependency of the UAS/Gal4 system, high expression levels of candidate genes were induced in adult flies starting at 24 h before injury. Axonal regrowth was defined as the growth of novel sprouts from the site of injury within four days. We used three parameters to evaluate axonal regrowth following injury: capacity of regrowth (the percentage of brains that exhibited at least one axonal sprout grown de novo), total regrowth (defined as the sum of the lengths of novel sprouts), and the maximum projection distance (defined as the distance of the longest novel sprout from the site of injury to their terminus) (Fig. 2 a-d). Of the genes tested, 7 (*drl*, *Dscam1*, *faf*, *kay*, *pdm2*, *pum* and *sens*) showed enhanced regeneration in all three categories (Fig. 2e-h). Kayak is the fly homologue of Fos, a key transcription factor downstream of JNK signaling, confirming that the screen can identify bona fide regeneration genes. Dichaete, D, is an example of a gene that promoted axonal outgrowth during development (Fig 2. a and d), but failed to induce regeneration in most cases (Fig. 3 a and d) and often resulted in short sprouts with poor morphology, making it difficult to measure. Three genes (*dimm*, d*ac* and s*qz*) caused axonal phenotypes such as defasciculation, blebbing or fragmentation, and were excluded from further analysis.

**Figure 1:**
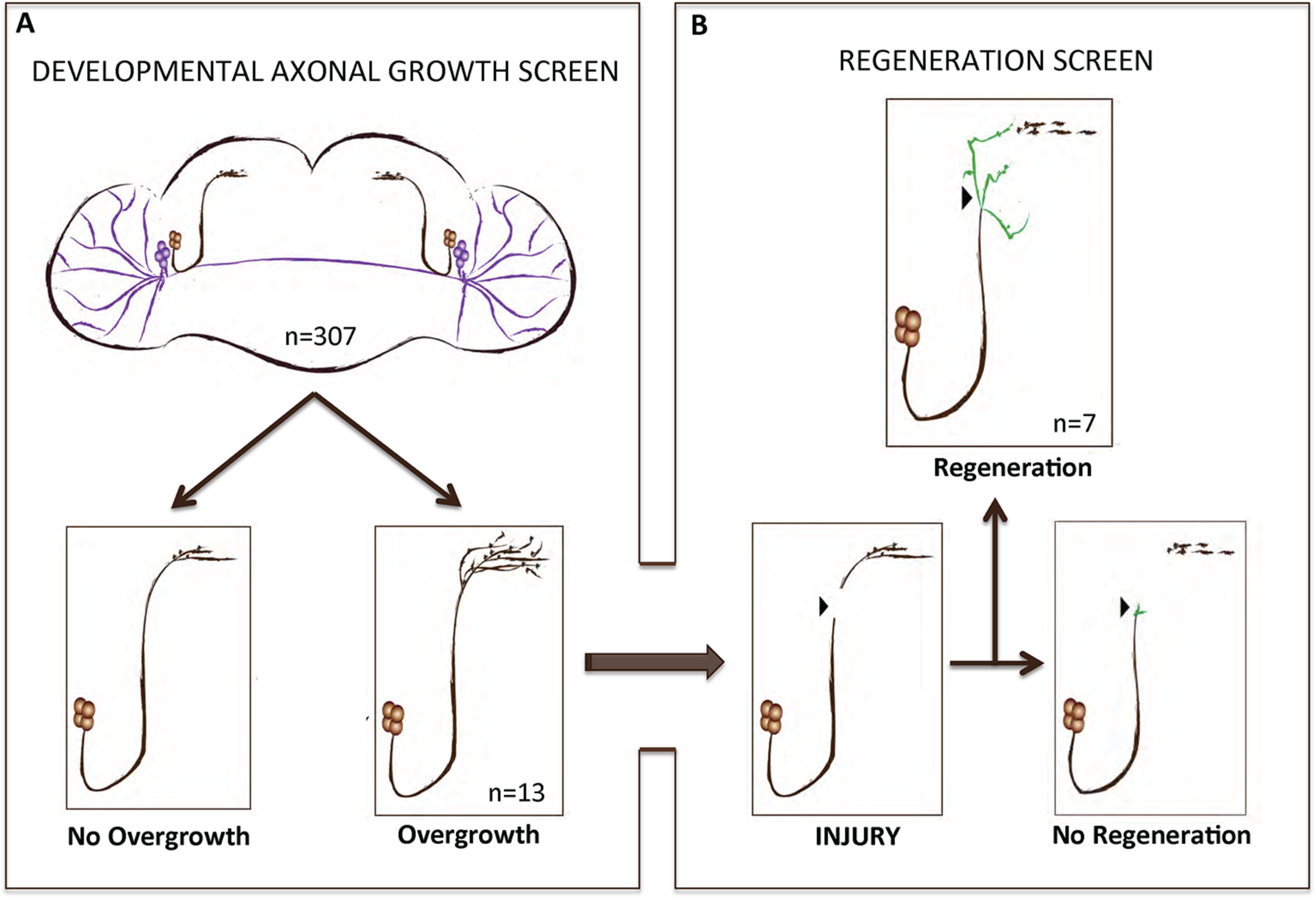
A gain of function screen for axonal growth in development and after injury. **(A,B)** Schematic illustrating the various outcomes for the development and injury steps of the screen. Both sLNVs (dark brown) and lLNVs (purple) are depicted in (A). Gain of function the candidate genes specifically in PDF neurons appeared to increase growth in 4.2% (n=13) of the cases. Genes that stimulated axonal growth were tested further in an injury paradigm in which the sLNV axonal projection is physically cut and sLNVs were accessed for regrowth four days post-injury (B). Seven genes retained the ability to promote significant regeneration of injured axons.

**Figure 2:**
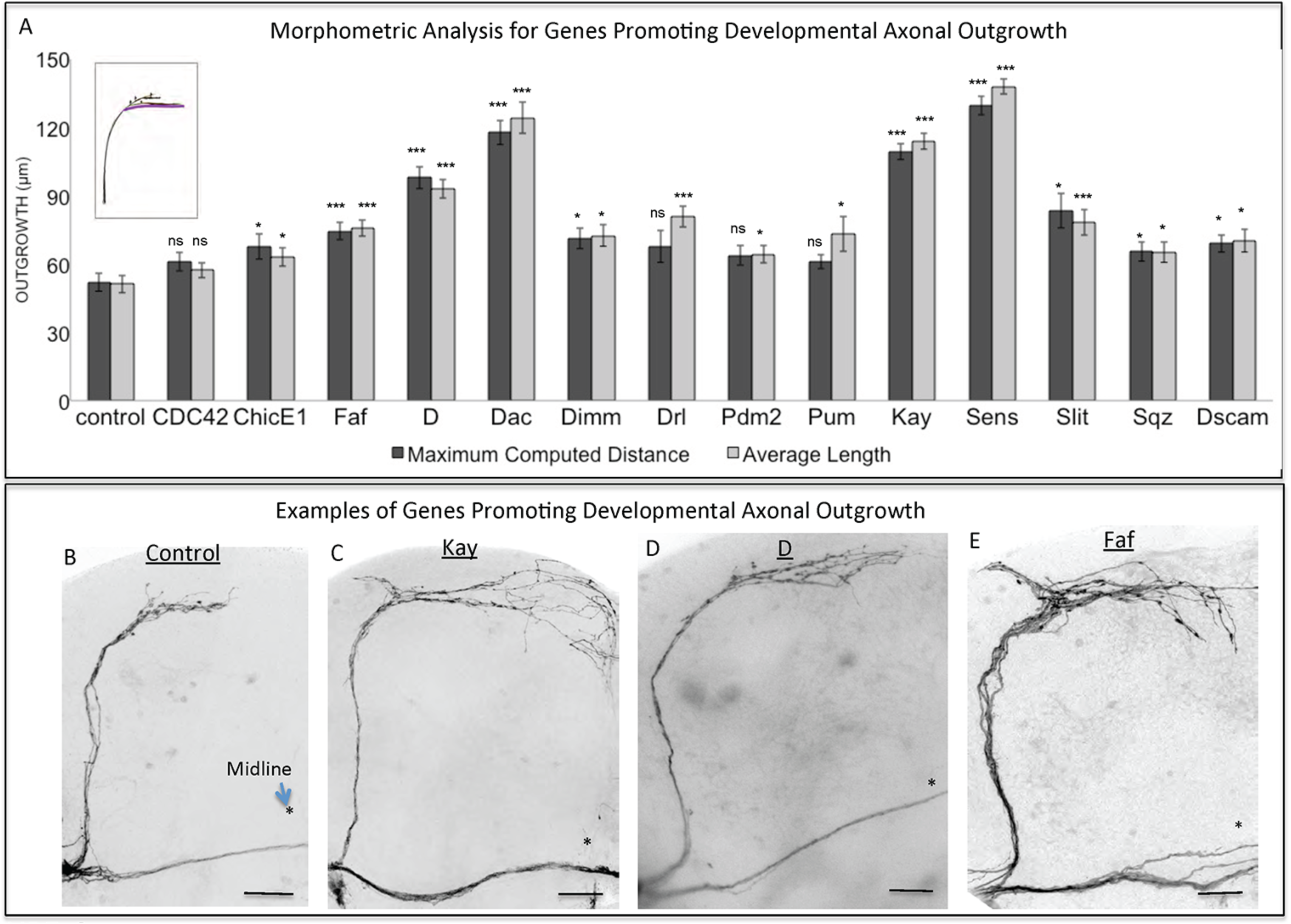
Analysis of axonal outgrowth in the developmental screen. **(A)** Morphometric analysis (Maximum Computed Distance and Average Length) of sLNv axonal projections where developmental overexpression of candidate genes has been specifically induced in the sLNvs. Axonal outgrowth is measured in μm. Purple trace in schematic represents measured axonal length. **(B-E)** Representative images of sLNv axonal arborization in wild type (control) adult flies (B) and in flies where developmental overexpression of *Kayak*, *kay* (C), *Dichaete*, *D* (D) and *Fat Facets*, *faf* (E) has been specifically induced in the sLNvs. Scale bars are 20 μm. Genotype of flies in (B) is PDF-Gal4, UAS-GFP/+; PDF-Gal4, UAS-2x eGFP/+, in (C) is PDF-Gal4, UAS-GFP/+; PDF-Gal4, UAS-2x eGFP/+; EP Kay/+, in (D) is PDF-Gal4, UAS-GFP/+; PDF-Gal4, UAS-2x eGFP/+; UAS-D/+, in (E) is PDFGal4, UAS-GFP/+; PDF-Gal4, UAS-2x eGFP/+; UAS-Faf/+. Asterisk denotes the brain midline, red arrow denotes the injury point. *, p < 0.05; ***, p < 0.001. n.s. indicates no statistical significance. Error bars represent SEM. Dotted insets have been zoomed in to better illustrate the diverse axonal phenotypes obtained. Scale bars are 20 μm.

### The ability of Faf to induce axonal regrowth is conserved and depends on its enzymatic activity

Of the identified 7 genes, 3 in particular (*Dscam1*, *faf* and *kay*) appeared to consistently promote robust growth. We therefore asked whether these genes might be acting together in a novel regeneration pathway linking the cell surface to the nucleus. We began by analyzing the de-ubiquitinating enzyme Faf since it promoted the highest levels of regeneration across all criteria (Fig. 3 a-c and h). First, we confirmed that *faf* is also able to induce axonal overgrowth in other CNS neuronal populations, such as the Dorsal Cluster Neurons (DCNs) (Figure 3 - Figure Supplement 1), suggesting that Faf may be a general CNS axonal growth-promoting factor. Ubiquitin-dependent protein regulation is critical in regulating many neuronal events, including axonal growth (Ambrozkiewicz and Kawabe, 2015; McCabe et al., 2004). However, the signaling pathways operating downstream of these enzymes are still largely unknown. To test whether the axonal growth induced by Faf was dependent on its deubiquitinase activity, we mutated a critical cysteine 1677 residue in the catalytic protease site to a serine (Chen and Fischer, 2000). In contrast to wild-type Faf, this mutated form of Faf was not able to significantly promote developmental axonal growth (Fig. 4 a,b and d). The mouse homologue of Faf, FAM/Usp9x, which can be active in *Drosophila* in other contexts (Chen et al., 2000; Wood et al., 1997), also induced robust sLNv axonal outgrowth (Fig. 4 c and d). Remarkably, even the yeast homologue of Faf, Ubp2, which only shares homology in the de-ubiquitination domain, induces sLNv axonal outgrowth very similar to Faf (Figure 4 – Figure Supplement 1). More importantly, both FAM and Faf, but not the enzymatic mutant Faf-Ser, induced significant axonal regeneration after injury (Fig. 3 e-h). These data suggest a conserved axonal growth and regeneration activity for Faf as a deubiquitinase enzyme.

**Figure 3:**
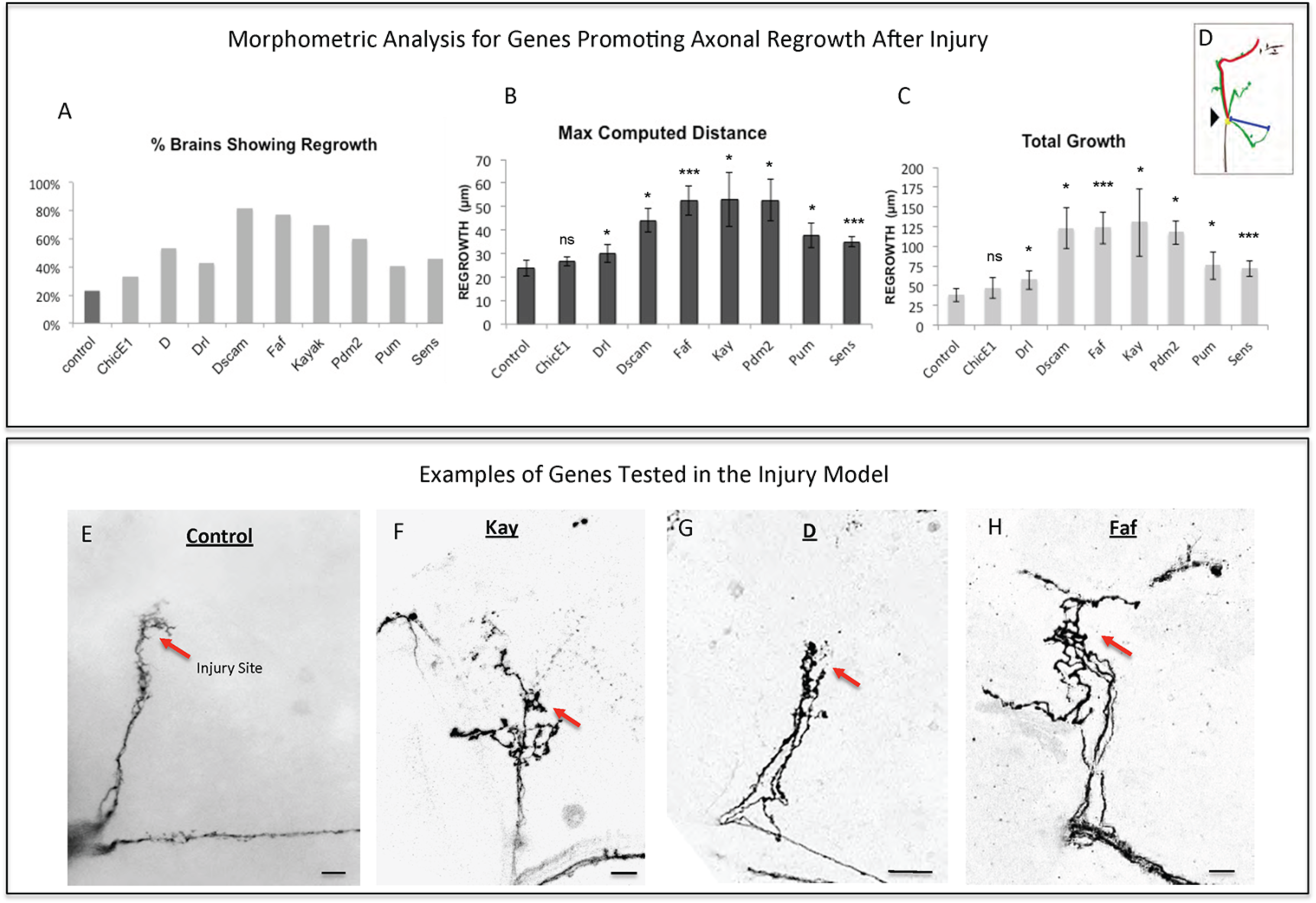
Analysis of axonal regrowth in the regeneration screen. **(A-D)** Analysis of axonal regrowth four days after injury. **(A)** Percentage of brains where at least one regenerated axonal sprout is detected (Capacity of regrowth) **(B,C)** Morphometric analysis (Maximum Computed Distance, (B) and Total Growth (C)) of regenerated sLNv axonal sprouts. Axonal regrowth is measured in μm. **(D)** Schematic simplifying how length of the regrown axonal sprouts is assessed. Yellow dot shows the point of injury; red trace represents maximum axonal length; blue trace represents axonal length measured in a straight line. **(E-H)** Representative images of sLNv axonal regrowth four days after injury in wild type (control) adult flies (E) and in flies where overexpression of *Kayak*, *kay* (F), *Dichaete*, *D* (G) and *Fat Facets*, *faf* (I) has been specifically induced in the sLNvs. Genotype of flies in (E) is PDF-Gal4, UAS-GFP/+; PDF-Gal4, UAS-2x eGFP/+, in (F) is PDF-Gal4, UAS-GFP/+; PDF-Gal4, UAS-2x eGFP/+; EP Kay/+, in (G) is PDF-Gal4, UAS-GFP/+; PDF-Gal4, UAS-2x eGFP/+; UAS-D/+, in (H) is PDFGal4, UAS-GFP/+; PDF-Gal4, UAS-2x eGFP/+; UAS-Faf/+. Asterisk denotes the brain midline, red arrow denotes the injury point. *, p < 0.05; ***, p < 0.001. n.s. indicates no statistical significance. Error bars represent SEM. Dotted insets have been zoomed in to better illustrate the diverse axonal phenotypes obtained. Scale bars are 20 μm.

**Figure 4:**
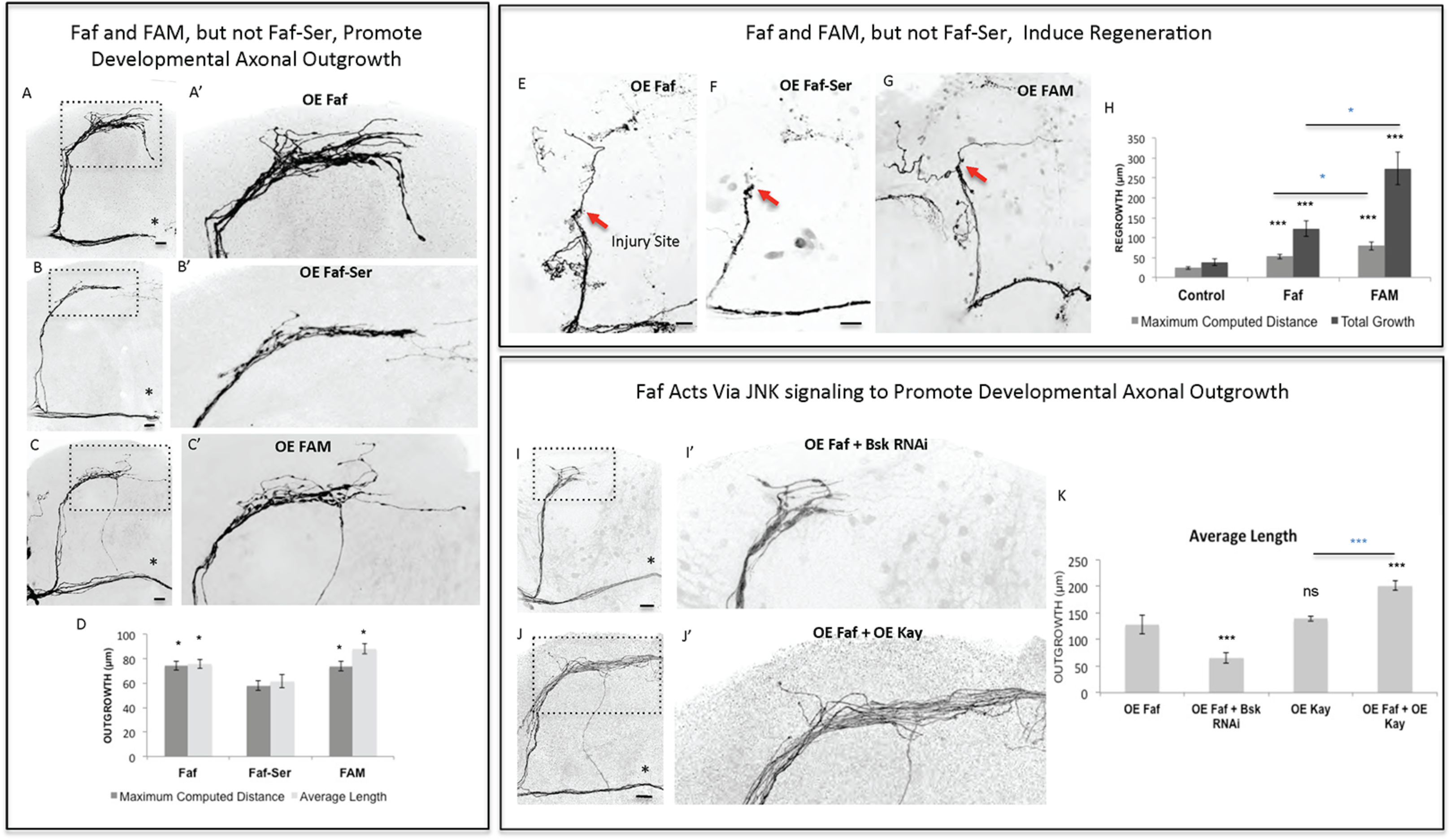
Faf and FAM, but not Faf-Ser, promote axonal outgrowth in development and axonal regrowth after injury, and interact with the JNK signaling pathway. **(A-C)** Representative images of sLNv axonal arborization in adult flies where developmental overexpression of *Fat facets*, *faf* (A and A’), *Fat-Serine*, *Faf-Ser* (B and B’), and *FAM* (C and C’) has been specifically induced in the sLNvs. **(D)** Morphometric analysis (Maximum Computed Distance and Average Length) of sLNv axonal projections for (A-C). Axonal outgrowth is measured in μm. **(E-G)** Representative images of sLNv axonal regrowth four days after injury in flies where overexpression of *faf* (E), *Faf-Ser* (F) and *FAM* (G) has been specifically induced in the sLNvs. **(H)** Morphometric analysis (Maximum Computed Distance and Total growth) of regenerated sLNv axonal projections in (E-G). Axonal outgrowth is measured in μm. **(I,J)** Representative images of sLNv axonal arborization for epistasis experiments between *faf* and *Bsk* (I and I’) and *faf* and *kay* (J and J’). **(K)** Morphometric analysis (Average Length) of sLNv axonal projections where developmental overexpression of *faf*; *faf* and *Bsk* RNAi; *kay*; and *faf* and *kay*, has been specifically induced in the sLNvs. Axonal outgrowth is measured in μm. Genotype of flies in (A, A’ and E) is PDF-Gal4, UAS-GFP/+; PDF-Gal4, UAS-2x eGFP/+; UAS-Faf/+, in (B, B’ and F) is PDF-Gal4, UAS-GFP/+; PDF-Gal4, UAS-2x eGFP/+; UAS-Faf-Ser/+, in (C, C’ and G) is PDF-Gal4, UAS-GFP/+; PDFGal4, UAS-2x eGFP/+; UAS-FAM/+, in (I and I’) is PDF-Gal4, UAS-GFP/+; PDFGal4, UAS-2x eGFP/+; UAS-Faf/UAS-Bsk RNAi, in (J and J’) is PDF-Gal4, UASGFP/+; PDF-Gal4, UAS-2x eGFP/+; EP-Faf/EP-kay. Dotted insets have been zoomed in to better illustrate the diverse axonal phenotypes obtained. Asterisk denotes the brain midline, red arrow denotes the injury point. *, p < 0.05; ***, p < 0.001. n.s. indicates no statistical significance. Error bars represent SEM. Dotted insets have been zoomed in to better illustrate the diverse axonal phenotypes obtained. Scale bars are 20 μm.

### Faf promotes axon regrowth in a JNK-dependent manner

Faf has been shown to induce neuromuscular junction growth in *Drosophila* (DiAntonio et al., 2001) in a pathway that requires Wallenda (Wnd), a conserved MAPKK upstream of JNK signaling (Collins et al., 2006). Therefore, we tested whether Faf required Wnd to induce axonal growth. RNA interference knock-down (RNAi KD) of *wnd* inhibited Faf-mediated axonal outgrowth (Figure 4 – Figure Supplement 2 a,b and i), whereas overexpression of *wnd*, but not a kinase-dead form of it, strongly promoted axonal outgrowth that essentially phenocopied *faf* overexpression (Figure 4 – Figure Supplement 2 a,c and d). Moreover, overexpression of *wnd* also promoted axonal regrowth after injury (Figure 4 – Figure Supplement 2 g,h and j). Therefore, Wnd likely acts downstream of Faf, to modulate axonal growth and regeneration in response to *faf* overexpression. Similarly, RNAi KD of the *Drosophila* homologue of JNK, *basket* (*bsk*), completely inhibited *faf*-mediated axonal outgrowth (Fig. 4. I and j). Conversely, co-expression of *kay*, the JNK pathway effector we identified as strong promoter of outgrowth in development (Fig. 2 a and c) and after injury (Fig. 3 a,b,c and f) enhanced Faf-mediated axonal outgrowth (Fig. 4 j and k). Together, these data suggest that Wnd and JNK act downstream of Faf to induce axonal outgrowth and regeneration.

### Faf stabilizes Dscam1 protein levels to promote axonal growth

How might *faf* activate JNK signaling to induce axonal regeneration? During fly eye development *faf* mediates the internalization of the Notch ligand Delta (Overstreet et al., 2004), and Notch signaling has been proposed to enhance regeneration of developing neurons (Kato et al., 2011), though it has also been shown to act as a repressor of axonal regeneration (El Bejjani and Hammarlund, 2012). To test if *faf* interacted with Delta in the context of sLNv axonal growth, we tested both a RNAi KD as well as a dominant negative (DN) transgene, and found that loss of Delta function in the sLNvs did not reduce the axonal outgrowth activity of *faf* (data not shown), suggesting an alternative mechanism in the context of axonal growth. Therefore, we reasoned that Faf might interact with different axon growth-promoting effectors.

Mammalian *Dscam1* (Qu et al., 2013) has been shown to be a regulator of JNK signaling. Interestingly, our screen identified *Dscam1* as one of the genes that most strongly and consistently promoted sLNv axonal outgrowth and regeneration (Fig. 2 a, Fig. 3 a-c). This prompted us to investigate the molecular mechanisms underlying the growth and regeneration activity of *Dscam1*. The *Dscam1* gene generates a large number of isoforms by alternative splicing of a plethora of extracellular domains and two transmembrane domains called TM1 and TM2 (Schmucker et al., 2000). We find that different isoforms containing either the TM1 (Fig. 2 a, Fig. 5 a), or the TM2 domain (not shown), and different extracellular domains (see methods) can induce axonal outgrowth, suggesting that induction of axonal growth may be a general property of Dscam1-mediated signaling independent of its isoform specificity. It has previously been reported that isoforms containing TM1 are dendrite specific (Shi et al., 2007). However, we find that upon overexpression these isoforms localize to both cell bodies and axonal terminals (Figure 5 – Figure Supplement 1). Conversely, *Dscam1* knockdown with two different RNAi lines (Watson et al., 2005) resulted in stunted sLNv axonal growth (Fig. 5 b and f). Finally, TM1-containing Dscam1 isoforms, UAS-Dscam1 1.30.30.1 GFP (Fig. 1 j-l) and UAS-Dscam1 1.34.31.1. HA induce robust axonal regeneration after injury, with the latter being the strongest line (Fig. 5 c).

**Figure 5:**
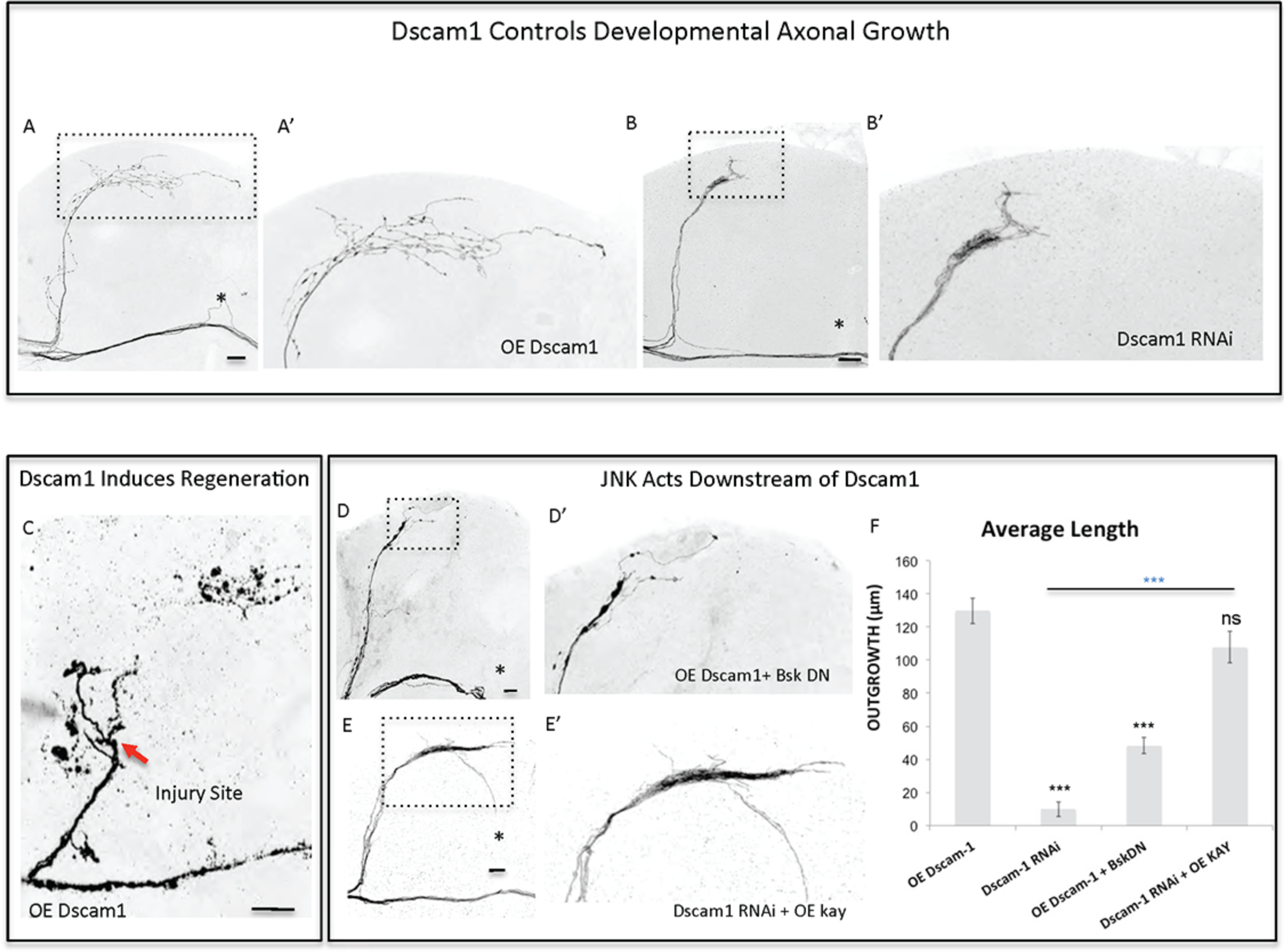
Dscam1 promotes axonal outgrowth in development and axonal regrowth after injury, and interacts with the JNK signaling pathway. **(A,B)** Representative images of sLNv axonal arborization in adult flies where developmental overexpression of *Dscam1* (A and A’), and Dscam1 RNAi (B and B’) has been specifically induced in the sLNvs. **(C)** Representative image of sLNv axonal regrowth four days after injury in flies where overexpression of *Dscam1* has been specifically induced in the sLNvs. **(D,E)** Representative images of sLNv axonal arborization demonstrating that inhibition of *Bsk* accomplished by overexpression of a dominant negative line of *Bsk* inhibits Dscam1-induced outgrowth (D and D’), and that overexpression of *kay* rescues the lack of axonal growth induced by overexpression of Dscam1-RNAi (E and E’). **(F)** Morphometric analysis (Average Length) of sLNv axonal projections where developmental overexpression of *Dscam1*; Dscam1-RNAi; *Dscam1* and Bsk DN; Dscam1-RNAi and *kay*, has been specifically induced in the sLNvs. Axonal outgrowth is measured in μm. Genotype of flies in (A,A’ and C) is PDF-Gal4, UAS-GFP/+; PDF-Gal4, UAS-2x eGFP/+; UAS-Dscam1-HA/+, in (B and B’) is PDF-Gal4, UAS-GFP/+; PDF-Gal4, UAS-2x eGFP/+; UAS-Dscam1-RNAi/+, in (D and D’) is PDF-Gal4, UAS-GFP/+; PDF-Gal4, UAS-2x eGFP/UAS-Bsk-DN; UAS-Dscam1-HA/+, in (E and E’) is PDF-Gal4, UAS-GFP/+; PDF-Gal4, UAS-2x eGFP/+; UAS-Dscam1-RNAi /EPkay. Dotted insets have been zoomed in to better illustrate the diverse axonal phenotypes obtained. Asterisk denotes the brain midline, red arrow denotes the injury point. *, p < 0.05; ***, p < 0.001. n.s. indicates no statistical significance. Error bars represent SEM. Dotted insets have been zoomed in to better illustrate the diverse axonal phenotypes obtained. Scale bars are 20 μm, with exception of C, which is 30 μm.

Both mammalian and fly Dscam1 are known to interact with p21 activating kinase (Pak) (Li and Guan, 2004; Schmucker et al., 2000) itself an upstream JNK Kinase. We find that inhibition of JNK activity, by using a dominant negative form of Bsk, completely abrogates Dscam1 mediated axonal growth (Fig. 5 d and f). Conversely, the expression of Kay completely reverses the loss of axon growth caused by *Dscam1* RNAi knock-down (Fig. 5 e and f). These data suggest that Dscam1 acts upstream of JNK signaling to induce axonal growth.

The fact that Faf and Dscam1 both promote axonal regeneration after injury and induce strikingly similar JNK-dependent axonal outgrowth phenotypes lead us to hypothesize that Faf and Dscam1 interact in this context. Indeed, we find that Faf-induced axonal outgrowth required Dscam1, as *Dscam1* knock-down almost completely abolished Faf-induced growth (Fig. 6 a and d). Consistent with this, co-overexpression of *faf* and *Dscam1* in the sLNvs induces stronger axonal outgrowth than *faf* overexpression alone (Fig. 6 b and d). Importantly, *Dscam1* knock-down also inhibits FAM/Usp9x mediated axonal outgrowth, indicating a conserved interaction (Fig. 6 c,d).

**Figure 6:**
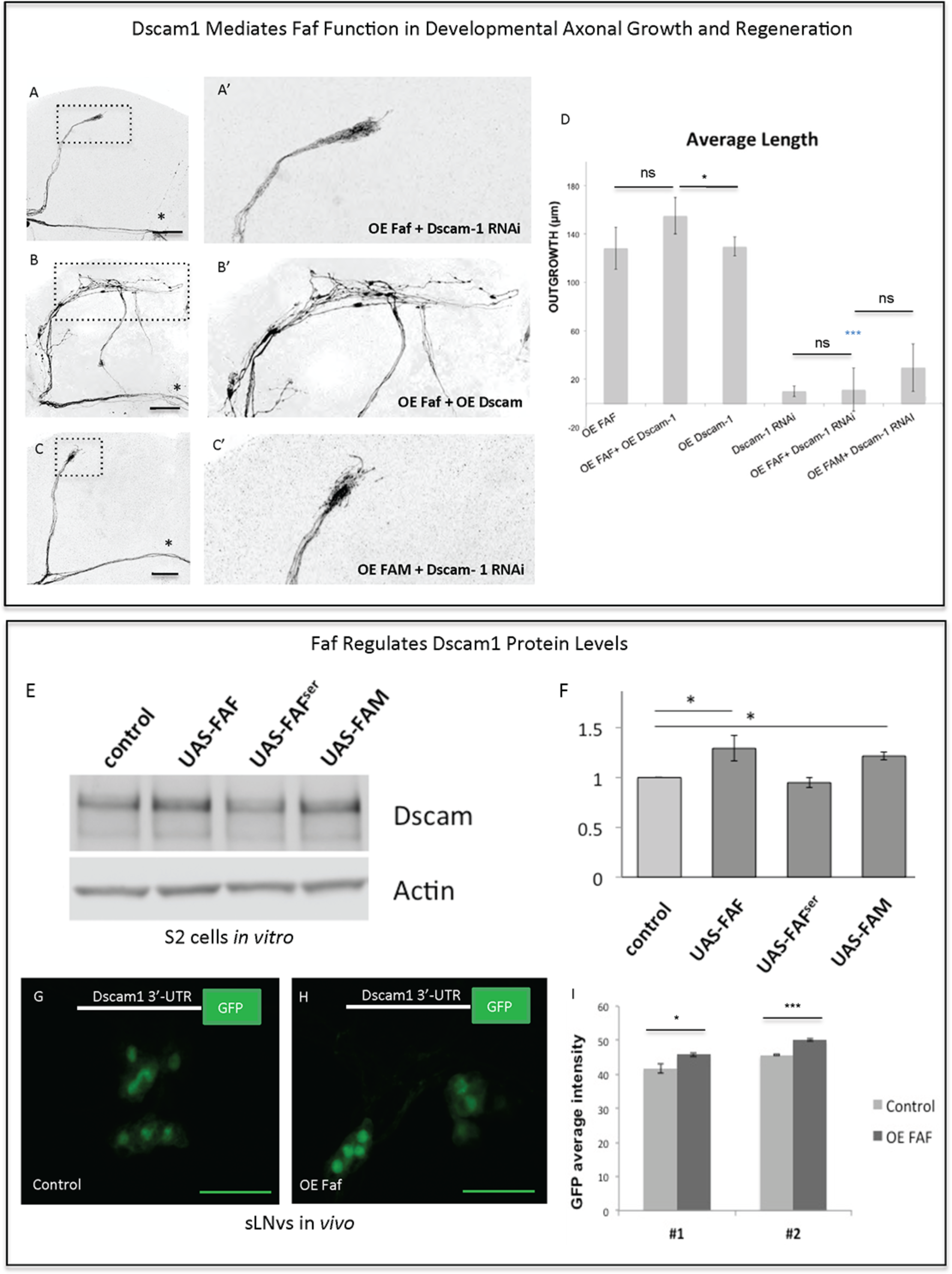
Faf and Dscam genetically and biochemically interact. **(A-C)** Representative images of sLNv axonal arborization demonstrating that knock-down of *Dscam1* inhibits Faf-induced outgrowth (A and A’), that cooverexpression of both *faf* and *kay* potentiates axonal growth (B and B’) and that knock-down of *Dscam1* inhibits FAM induced outgrowth. **(D)** Morphometric analysis (Average Length) of sLNv axonal projections where developmental overexpression of *faf*, *FAM*, *Dscam1* and Dscam1 RNAi has been specifically induced in the sLNvs, uncovering gene interactions. Axonal outgrowth is measured in μm. **(E,F)** Western blot and quantification showing increased levels of Dscam1 protein following S2 electroporation of wild-type Faf and FAM, but not of Faf-Ser in comparison to control (UAS vector). **(G-I)** GFP fluorescence analysis showing increased levels of 3’UTR-Dscam1 following overexpression of *faf* (n=28) (H) in comparison to its control (n=24) (G). LNVs GFP average intensities are shown in (I). Genotype of flies in (A and A’) is PDF-Gal4, UAS-GFP/+; PDF-Gal4, UAS-2x eGFP/+; / UAS-Faf/UAS-Dscam1-RNAi, in (B and B’) is PDF-Gal4, UAS-GFP/+; PDF-Gal4, UAS-2x eGFP/+; / UAS-Faf/UAS-Dscam1-HA, in (C and C’) is PDFGal4, UAS-GFP/+; PDF-Gal4, UAS-2x eGFP/+; UAS-FAM/ UAS-Dscam1-RNAi, in (G) is PDF-Gal4, UAS-GFP/+; UAS-3’-UTR-Dscam1-GFP/+; UAS-Faf/+, in (H) is PDF-Gal4, UAS-GFP/+; UAS-3’-UTR-Dscam1-GFP/+; TM6b/+. Dotted insets have been zoomed in to better illustrate the diverse axonal phenotypes obtained. Asterisk denotes the brain midline, *, p < 0.05; ***, p < 0.001. n.s. indicates no statistical significance in (D). Error bars represent SEM in (D) and in (I). Scale bars in (A-C) and (G-H) are 30 μm.

Faf antagonizes ubiquitination by cleaving the covalent bond between ubiquitin and a substrate protein (Huang et al., 1995), thereby leading to stabilization of proteins targeted for degradation. We asked if Faf might stabilize Dscam1 protein levels. Therefore, we expressed *Dscam1* alone or together with *faf, faf-Ser* mutant or mouse *FAM/Usp9x* in *Drosophila* S2 cells. Both Faf and FAM/Usp9x, but not the Faf-Ser mutant lead to a ~30% increase in Dscam1 protein levels (Fig. 6 e and f), with no change in mRNA levels (Supplementary data file 2). However, we were unable to find evidence for Dscam1 ubiquitination in wild type or proteasome-inhibited S2 cells, nor a change in that status upon overexpression or knock-down of *faf* (data not shown). These data suggest that, at least in this context, Faf does not de-ubiquitinate Dscam1 directly. We then asked whether Faf enhances translation of Dscam1 through a mechanism that is mediated by the 3’UTR of *Dscam1*. To this end, we expressed a *Dscam1-3’-UTR>GFP* reporter in the sLNv alone or together with *faf*. We find that GFP levels are significantly upregulated in sLNv upon Faf overexpression (Fig 6 g,h and i).

Interestingly, Wnd has been shown to be a positive upstream regulator of Dscam1 in neurons. Specifically, using the same *Dscam1-3’-UTR>GFP* reporter construct it has been shown that translation of the reporter protein is enhanced by Wnd (Kim et al., 2013). Therefore, we tested whether Dscam1 is also required for Wnd-induced axonal growth in sLNv neurons and find that Dscam1 RNAi KD significantly decreases Wnd-induced growth (Figure 4 – Figure Supplement 2 e and f). Finally, we asked whether Faf enhances translation of Dscam1 through a mechanism that is mediated by the 3*’*UTR of *Dscam1*. To this end, we expressed the *Dscam1-3’-UTR>GFP* reporter in the sLNv alone or together with *faf*. We find that GFP levels are significantly upregulated in sLNv upon Faf overexpression (Fig 6 g,h and i).

## Discussion

In contrast to young neurons, injured adult CNS neurons exhibit very limited ability to self-repair, suggesting that the intrinsic regenerative capacity is lost during development. For example, it has been shown that the axon growth rate decreases dramatically with age in post-natal retinal ganglion cells (Goldberg et al., 2002). In addition, pioneer work from Filbin and colleagues demonstrated that developmental loss of the regenerative capacity of neurons in post-natal rats is mediated by a decline in the endogenous levels of neuronal cAMP within a few days after birth (Cai et al., 2001). Consistent with this evidence, transcription factors that regulate developmental axonal growth, such as members of the Kruppel-like family (KLFs), can promote regrowth of adult injured corticospinal tract and optic nerve axons (Blackmore et al., 2012; Moore et al., 2009). Several other intrinsic axonal regulators, including phosphatase and tensin homolog (PTEN), suppressor of cytokine signaling 3 (SOCS3), mTOR, Osteopontin and IGF-1 (Duan et al., 2015; Liu et al., 2010; Sun et al., 2011), have been previously identified in mammalian systems-though mostly on a gene by gene basis. More systematic approaches, such as quantitative proteomic analysis, have been recently employed to identify molecular pathways that are altered in injured retinal ganglion cells, and identified additional intrinsic regulators of regeneration, such as c-Myc (Belin et al., 2015). Taken together, all these studies suggest that manipulation of the intrinsic regenerative ability of mature neurons might be efficient strategies for enhancing the capacity of injured axons to regenerate. It is therefore crucial to discover factors that constitute intrinsic pro-regeneration signaling pathways in a systematic manner. We have previously shown that the adult *Drosophila* CNS is a suitable model for studying axonal injury and regeneration (Ayaz et al., 2008). By exploiting this model along with the power of *Drosophila* genetic screens, we have uncovered a novel axonal regeneration pathway (Fig. 7) that links the stability of the neuronal cell surface receptor Dscam1, via the de-ubiquitination function of the enzyme Faf, to JNK signal, a major inducer of axonal regeneration in *C. elegans, Drosophila* and mouse (Ayaz et al., 2008; Li et al., 2012; Nix et al., 2011; Raivich et al., 2004; Raivich and Makwana, 2007).

E3 ubiquitin ligases are important in many aspects of mammalian brain development and function, by controlling neuritogenesis, modulating axon guidance and pruning, neuronal polarity and synaptic transmission (Ambrozkiewicz and Kawabe, 2015). Not surprisingly, E3 ligase dysfunction and abnormal ubiquitin signaling is implicated in several human brain disorders. In particular, mutations in the mammalian homologue of Faf, FAM/Usp9x, have been associated with X-linked intellectual disability (Homan et al., 2014). Brain specific deletion of FAM/Usp9x results in early postnatal death, and FAM/Usp9x knock-out neurons display reduced axon growth and impaired neuronal migration (Homan et al., 2014; Stegeman et al., 2013).

Ubiquitin-dependent signals that include Faf have also been shown to regulate synaptic development and growth at the *Drosophila* neuromuscular junction (NMJ) (DiAntonio et al., 2001), though its function in the CNS remained elusive. Both yeast, Ubp2, and mouse homologues of Faf display the ability to induce axonal growth in the CNS, suggesting conservation of this property throughout evolution, similar to what had been shown at the NMJ (DiAntonio et al., 2001),(Kim et al., 2013). Importantly, we show for the first time that both Faf and FAM promote regrowth of injured axons in the adult fly brain.

Ubiquitin signaling is rather complex, and dissecting it downstream players can be challenging. In our screen, overexpression of the neuronal cell adhesion Dscam1 induced robust axonal growth both in development as well as after injury similar to the one induced by Faf, which led us to hypothesize that both genes acted in the same growth-promoting signaling pathway. In *Drosophila*, Dscam1 shows extensive molecular diversity that results from alternative splicing into some 18500 diverse extracellular domains (Schmucker et al., 2000). This isoform diversity has been shown to be critical for neuronal self-recognition and self-avoidance underlying axon growth and dendritic patterning (He et al., 2014; Hughes et al., 2007). Independent of its ectodomain diversity, Dscam1 has also been recently shown to regulate presynaptic arbor growth (Kim et al., 2013).

Our data indicate that post-transcriptional regulation of Dscam1 allows axonal growth after injury (Fig. 6). The Dscam1-stabilizing function of Faf appears to be conserved in mammals, as FAM/Usp9x overexpression also leads to increased levels of Dscam1 protein (Fig. 6 e,f). The placing of Faf upstream of Dscam1 and Wnd in regulating axonal outgrowth suggests that the de-ubiquitination activity of Faf may operate by antagonizing the E3 ubiquitin ligase activity of enzymes such as Highwire (DiAntonio et al., 2001). Interestingly, Wnd itself is a promoter of mRNA stability and local translation, and is essential for axon regeneration after laser axotomy in adult neurons in *C. elegans* (Byrne et al., 2014; Yan et al., 2009).

Remarkably, a screen for axonal injury response in the mouse identified the differential regulation of transcript-availability and -loading onto ribosomes of CNS development genes as a major feature of abortive regenerative response in the mammalian spinal cord (see accompanying manuscript). Furthermore, overexpression of a post-transcriptional regulator, the cytoplasmic polyadenylation element-binding protein1/Orb, in both *Drosophila* and mouse neurons is sufficient to induce axonal regrowth after injury. Together, our studies, also point to a crucial role of mRNA stability and translational control of neurodevelopmental genes in the design of future therapeutic strategies.

## METHODS

### Candidate gene sample

We used the Gene Ontology tool (<u>http://geneontology.org/)</u> to select candidate genes annotated with the terms ‘Neurite Morphogenesis’, ‘Transcription Factors’, ‘Receptors’, ‘Chromatin Modifiers’ and ‘Ubiquitin Ligases’. We also included an additional set of genes previously implicated in axonal growth and/or involved in actin dynamics (indicated in Supplementary data file 1 by asterisks).

The final candidate gene sample only included genes for which appropriate gain-of-function fly lines were readily available at the stock centers at the time of the study (Supplementary data file 1).

### Fly stocks and genetics

*Drosophila melanogaster* stocks were kept on standard cornmeal media. For tissue-specific overexpression of the transgenes, we used the GAL4/UAS system (Brand and Perrimon, 1993). Lines with UAS insertion sites (*i.e.* UAS, EP, EPgy2, XP and Mae-UAS) were received through the Bloomington or Szeged Stock Centres (or from specific laboratories when specified). Loss of function lines (Wnd RNAi GD8365, Dscam1 RNAi KK108835; Dscam1 RNAi (Watson et al., 2005); Bsk RNAi BL35594 and BL36643, and Bsk DN BL6409) were obtained from the Bloomington Stock Centre or from the Vienna *Drosophila* Research Centre (VDRC). UAS-Wnd kinase dead (KD) and UAS-Wnd E flies were a gift from C. Collins. The *PDF-Gal4* line was obtained from P. Taghert. For Faf overexpression in flies, we used the EP3520 line (Szeged Stock Centrum), which was previously reported to induce Faf gain-of-function (DiAntonio et al., 2001). UAS-Faf and UAS-FAM lines were created in house by cloning Faf cDNA and FAM cDNA into a pUAST-attB vector, respectively (Bischof et al., 2007) and injected in an attP2 docking line (BL 8622). A UAS-Faf serine (Faf-Ser) mutant that harbors a cysteine to serine mutation at residue 1677 was also cloned using the same method. UAS-Dscam1 HA-FLAG and UAS-Dscam1 GFP flies, both containing the 1.30.30.1 isoform, were used for the injury experiments. The UAS-Dscam1 1.30.30.1 GFP flies have been described (Hughes et al., 2007) and the UAS-Dscam1 HA-FLAG flies were created by inserting a HA tag into the intracellular domain (after the 81st bp of exon 22) of isoform 1.34.31.1. The UAS-Dscam1 1.30.30.1 GFP flies have been used for the initial screen in development and after injury, and the UAS-Dscam1 1.30.30.1 HA has been used for additional confirmatory experiments in development and injury, as well as for epistasis experiments. While both lines induce significant axonal growth in development and after injury, UAS-Dscam1 1.30.30.1 HA appears to be the strongest of the two lines. For the genetic screen in development and after injury, pdf-Gal4, UASGFP; pdf-Gal4, UAS-2x eGFP/cyo flies were kept as a stock and used to drive expression of the various candidate genes, or crossed to wild-type Canton S (CS) flies. For the genetic epistasis experiments, pdf-Gal4, UAS-GFP; UAS-Dscam1 RNAi and pdf-Gal4, UAS-GFP; Faf EP 3520 flies were maintained as a stock and crossed to overexpression lines to uncover genetic interactions.

### Developmental outgrowth screen

To measure axonal outgrowth during development, flies were reared at 25°C and were dissected 2-10 days after eclosion. A minimum of 5 fly brain (10 sLNs projections) per genotype were stained with an anti-GFP antibody (to enhance the GFP signal), visualized under a fluorescent microscope equipped with a GFP filter and scored as ‘growth’ (when sLNv axonal projections appeared considerably longer than in controls) or ‘no-growth’ (when the length of sLNv projections was indistinguishable from controls or shorter). All genes were scored growth promoting genes were confirmed as such in at least one independent experiment, and their growth inducing ability analysed by measuring the axonal sLNv dorsal axonal projections.

### Whole brain explant culture injury system

For the axonal regrowth analysis after injury, flies were reared at 18°C, in order to minimize overexpression effects during development, and shifted to 25°C the day before injury to allow optimal transgene expression.

Whole-brain explants on culture plate inserts were prepared and injured as described (Ayaz et al., 2008; Koch, 2012). In brief, Millicell low height culture plate inserts (Milipore) were coated with laminin and poly-lysine (BD Biosciences). Adult female flies were collected 2-10 days after eclosion and placed on ice. Fly brains were quickly and carefully dissected out in a sterile Petri dish containing ice cold Schneider’s *Drosophila* Medium (GIBCO). Up to seven brains were placed on the membrane of one culture plate insert and culture medium (10 000 U/ml penicillin, 10 mg/ml streptomycin, 10% Foetal Bovine Serum and 10 μg/ml insulin in Schneider’s *Drosophila* Medium) was added. sLNv axonal injury was performed using an ultrasonic microchisel controlled by a powered device (Eppendorf). Culture dishes were kept in a plastic box in a humidified incubator at 25°C

### Immunohistochemistry

Freshly dissected brains of adult flies were fixed in 4% formaldehyde and processed for immunohistochemistry as described (Hassan et al., 2000). Cultured brains (four days post-injury) were first fixed by replacing the culture medium in the Petri dish for 30 minutes. Then, 1 ml of fixative was carefully added on top of the filter for 1-2 hours. Brains that detached from the membrane were excluded from further analysis. Immunostaining was performed as for freshly dissected samples. Primary antibodies were rabbit anti-GFP (A-6455, Molecular Probes), rat anti-HA (3F10, Roche); and anti-Pdh (gift from P. Taghert).

### Imaging and morphological analysis of sLNv axonal projections in development and after injury

Image J software was used to measure the length of the dorsal axonal projections emanating from the sLNvs. The starting point was set as the point where axons turn medially and start to run parallel to the commissure. Axonal length was measured as a straight line (Computed Distance) from the starting point towards the midline (indicated by an asterisk) and as manual trace using Image J. The maximum computed distance was defined as the distance projected by the longest axonal sprout in a straight line and parallel to the commissure. The Average Length was the defined as the average length of the two longest axonal branches traced manually (freehand distance). Imaging was performed on an upright Zeiss Axioscope equipped with a CCD camera, or on a Zeiss 700 or Nikon AR1 confocal microscope. All measurements were performed using ImageJ.

To analyse the role of the candidate genes in axonal regrowth after injury we imaged cultured brains at two different time points after injury: approximately five hours and four days. Comparison between these two timepoints allowed us to define the location at which the injury took place, in order to define *de novo* growth. Morphometric analysis of axonal regrowth was always performed four days after injury, following fixation and GFP staining of the brains in culture. Capacity of regrowth was defined as the ability of the injured sLNv projection to regrow at least one new axonal sprout. Without the support of the head cuticle, brains will flatten and therefore undergo slight morphological changes during the culture process. To be conservative and account for potential inaccuracies in defining the injury point, only regrown axons with a minimum length of 12 μm were defined as de novo growth and taken into account for analysis. To quantify axonal regrowth, newly grown axons were measured in a straight line and manually traced using Image J. In this case, the maximum computed distance was defined as the average of the distance of the two longest axonal sprouts in a straight line in any direction. Maximum growth was defined as the sum of the freehand lengths of all de novo grown axons. Images five hours after injury were acquired on an upright Nikon microscope equipped with a Hamamatsu CCD camera ORCA-R2. Imaging 4 days after injury was performed on a Zeiss 700 or a Nikon A1R confocal microscope after GFP immunostaining. See figure legends for details of individual experiments, including statistical tests used and number of samples tested.

### Cell culture and western blotting

Drosophila Schneider’s (S2) cells were maintained in Sf-900 II SFM medium (Gibco). To achieve transgene overexpression in Schneider’s (S2) cells we electroporated a UAS construct in combination with PMT-Gal4, according to previously developed methods (Klueg et al., 2002). For Faf and FAM overexpression, we created a UAS-Faf and a UAS-FAM construct as described in “Fly stocks and Genetics”. For Dscam1 overexpression, the UAS-Dscam1 1.30.30.1 GFP construct was used (Hughes et al., 2007).

Cells were electroporated using an Amaxa Nucleofector KitV (Lonza), according to the manufacturer’s instructions. Cells were harvested 72-96 hours after copper induction, briefly washed with PBS and pellets frozen until cells were lysed in a 1% NP40 buffer in Tris-HCL. Protein concentration was determined by a modified Lowry assay (Peterson, 1977). Western blotting was performed with a SDSPAGE Electrophoresis System (Biorad). Briefly, protein samples were diluted in SDS containing sample buffer and 15 μg per sample was loaded onto a 3-8% Tris-Acetate mini gel (Novex, Life Technologies). Samples were blotted using tank transfer to a nitrocellulose membrane (GE Healthcare), blocked with milk and probed with primary antibodies against Dscam1 (1:1000) (Watson et al., 2005) or against Actin (1:5000, ab3280, Abcam), which was used as a protein loading control. Anti-rabbit or anti-mouse horseradish peroxidase conjugated secondary antibodies (Amersham) were then added, and proteins were detected using enhanced chemiluminescence (ECL Plus, GE Healthcare) on a FUJI LAS imager system (Fuji). Values for Dscam1 were normalized to the values of the loading control (actin) and quantified using the blot analysis function for IMAGE J. Kruskall Wallis test was used to compare the different conditions. Data is shown as mean ± SEM and significance was set at p≤0.05.

### RNA isolation and quantitative PCR

RNA was extracted with Trizol. 1 μg of total RNA was reverse transcribed using the Quantitect RT kit (Qiagen). qPCR using the Taqman Real Time protocol (Applied Biosystems) and probes (See Supplementary data file 2 for info Taqman probes). Data is shown as mean ± SEM in Supplementary data file 2.

### GFP intensity measurements

Adult brains were dissected and immediately prepared for imaging. Confocal stacks of all sLNv and lLNv cell bodies in each side of the brain were performed. The optimal confocal settings were first adjusted for wild type brains and kept unchanged to allow comparison between genotypes. A maximum projection was created for each brain side and each image was quantified for GFP intensity using the ‘Image Analysis’ module of Zeiss Zen 2.0 software. All quantifications were done by an investigator blind to experimental conditions. Student T test was used to compare both genotypes. Data is shown as mean ± SEM and significance was set at p≤0.5.

## SUPPLEMENTAL FIGURE LEGENDS

**Supplemental Figure 1:**
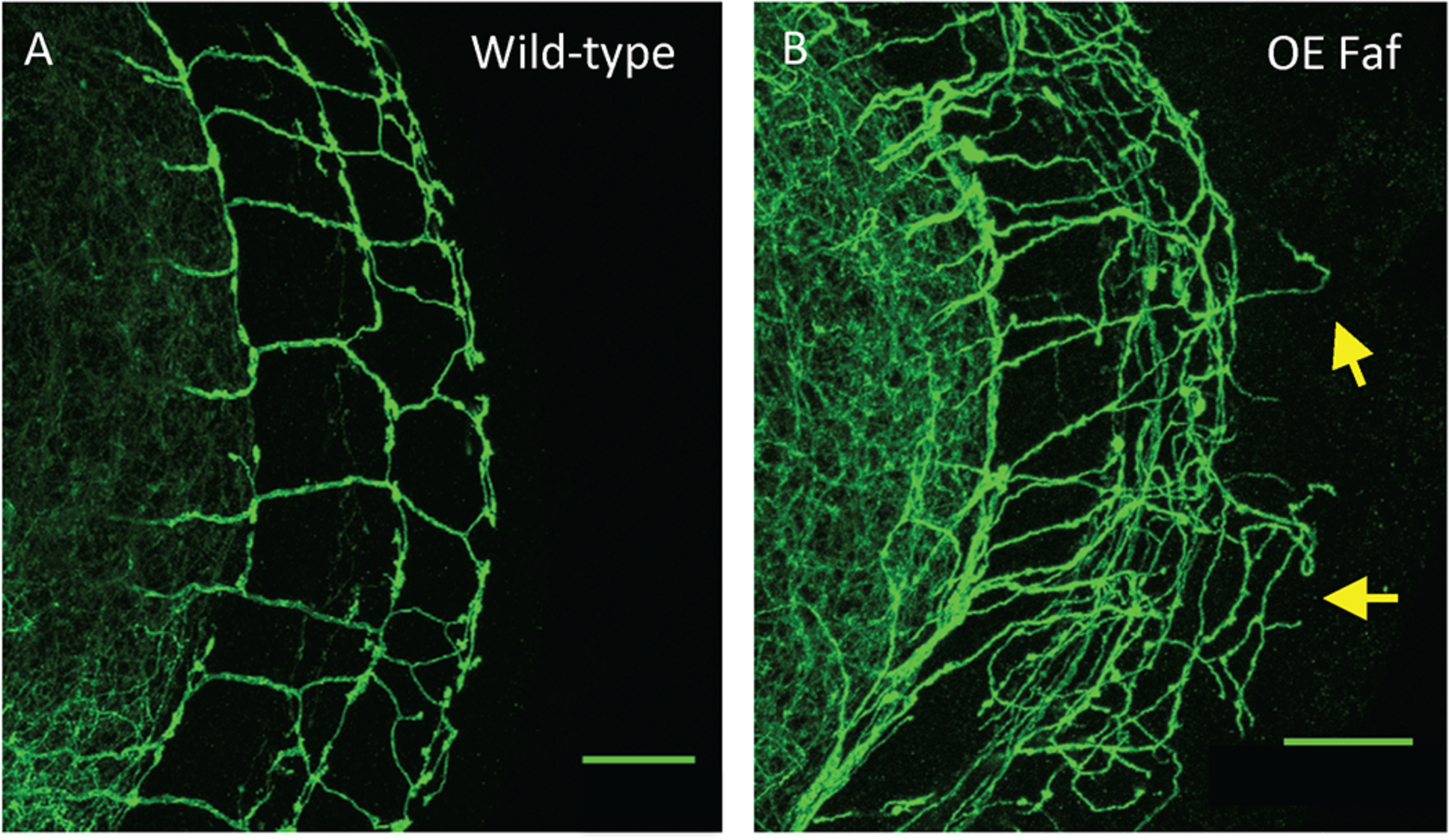
Faf gain of function promotes axonal growth in a distinct neuronal population. **(A,B)** Overexpression of *faf* specifically in the Dorsal Cluster Neurons (DCNs) results in increased axonal growth (yellow arrows) (B), in comparison to wild-type flies (A). Genotype of flies in (A and A’) is; UAS-GFP;ato-Gal4 14a, in (B and B’); UASGFP;ato-Gal4 14a/UAS-Faf. Scale bars are 20 μm.

**Supplemental Figure 2:**
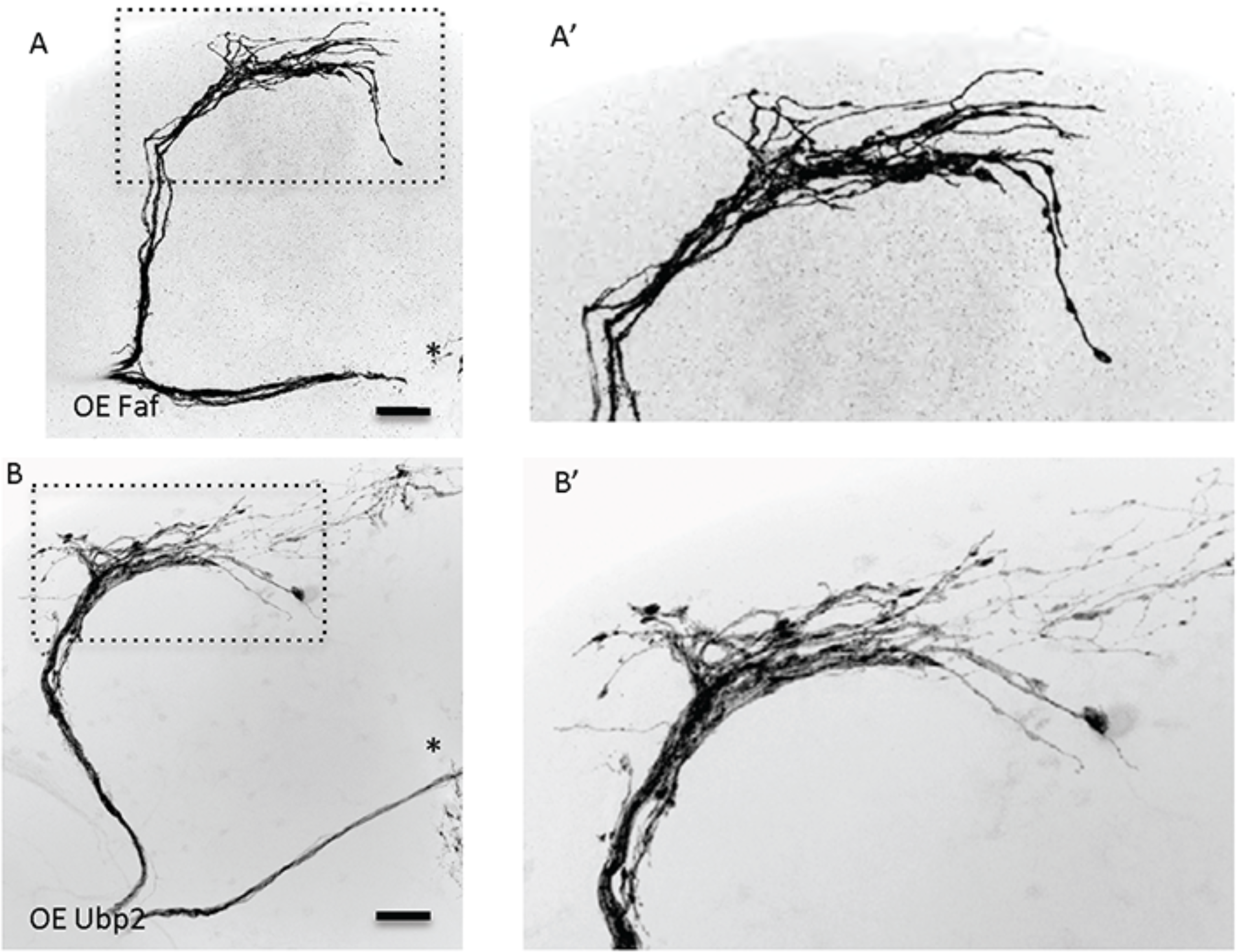
Ubp2 gain of function also promotes axonal growth. **(A,B)** Overexpression of the yeast homologue of Faf, *Ubp2*, which shows conservation of the enzymatic domain, also results in increased axonal growth (B), in a similar manner to Faf overexpression (A). Note that B and B’ are the same as in Fig. 4 A and A’ Genotype of flies in (A and A’) is PDF-Gal4, UAS-GFP/+; PDF-Gal4, UAS-2x eGFP/+; UAS-Faf/+; in (B and B’) is PDF-Gal4, UAS-GFP/+; PDF-Gal4, UAS-2x eGFP/UAS-Ubp2. Dotted insets have been zoomed in to better illustrate the diverse axonal phenotypes obtained. Scale bars are 30 μm.

**Supplemental Figure 3.**
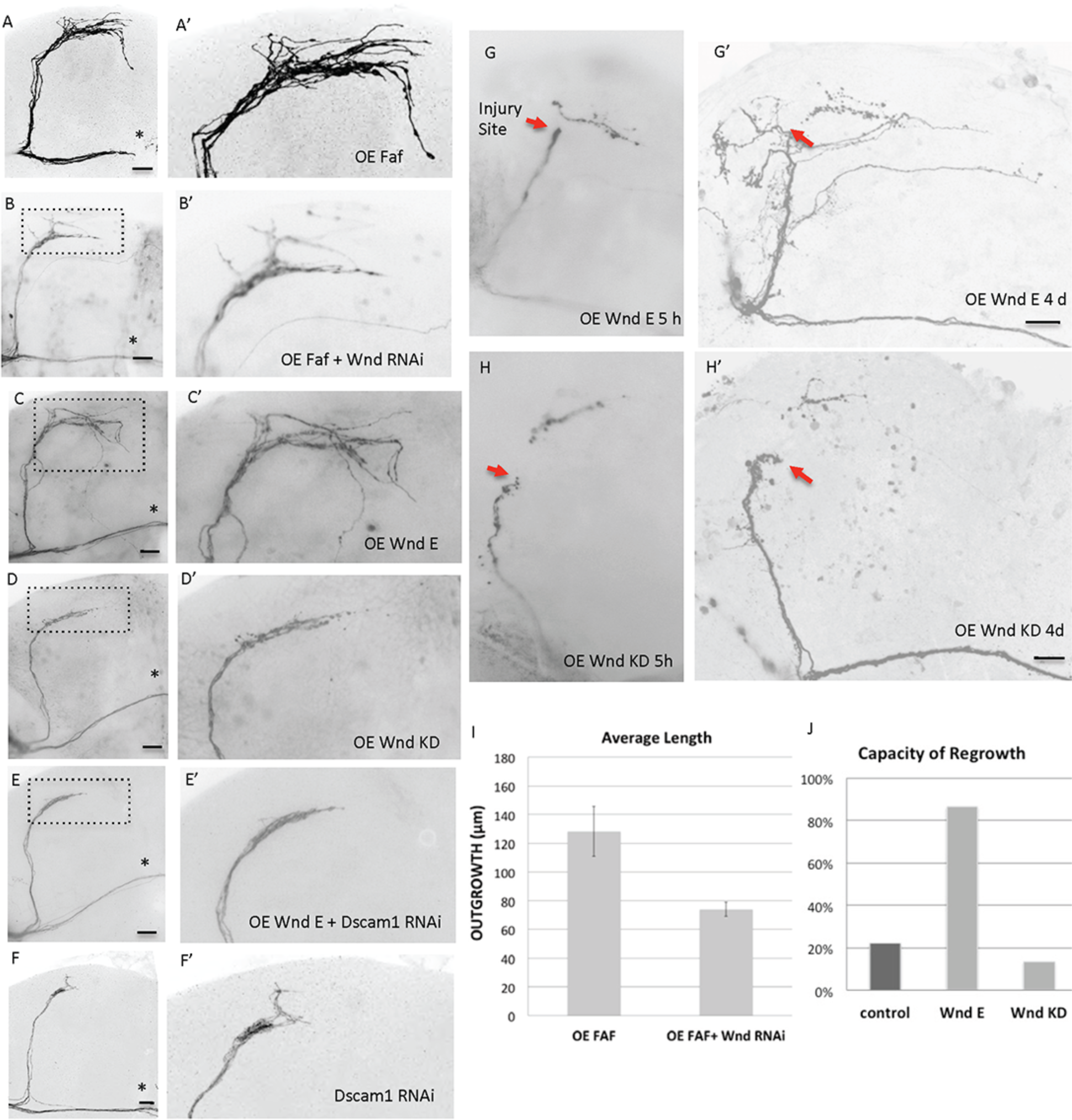
Wallenda promotes growth in development and after injury and is required for Faf-induced growth. **(A,B)** Representative images of sLNv axonal arborization demonstrating that knock-down of *wnd* inhibits Faf-induced outgrowth. **(C,D)** Representative images of sLNv axonal arborization in adult flies where developmental overexpression of *wnd* (C and C’), but not of a kinase dead form (D and D’) in the sLNvs induces axonal growth similar to the one induced by Faf (A and A’). **(E,F)** Representative images of sLNv axonal arborization demonstrating that knock-down of *Dscam1* inhibits Wnd-induced outgrowth (E and E’) and results in a phenotype that resembles knock-down of *Dscam1* on its own (F and F’). **(G,H)** Overexpression of *wnd* in the sLNVs (G and G’), but not of Wnd KD (H and H’) induces axonal regrowth four days after injury. Five hour after injury timepoints (G and H) have been included to better illustrate the regenerative ability of Wnd, but not of its kinase dead (KD) form. Red arrows point to the place of injury. **(I)** Morphometric analysis (Average Length) of sLNv axonal projections where developmental overexpression of *faf* and Wnd RNAi has been specifically induced in the sLNvs, uncovering a Faf-Wnd gene interaction. Axonal outgrowth is measured in μm. **(J)** Percentage of brains showing at least one regenerated axonal sprout four days after injury (Capacity of regrowth), where overexpression of *wnd* and Wnd KD has been specifically induced in the sLNvs. Note that A and A’ are the same as in Fig. 4 A and A’, and F and F’ the same as in Fig. 5 B and B’ Genotype of flies in (A and A’) is PDF-Gal4, UAS-GFP/+; PDF-Gal4, UAS-2x eGFP/+; UAS-Faf/+, in (B and B’) is PDF-Gal4, UAS-GFP/+; PDF-Gal4, UAS-2x eGFP/+; UAS-Faf/Wnd RNAi;, in (C, C’ and G,G’) is PDF-Gal4, UAS-GFP/+; PDF-Gal4, UAS-2x eGFP/UAS-Wnd E;, in (D,D’ and H,H’) is PDF-Gal4, UASGFP/+; PDF-Gal4, UAS-2x eGFP/+; UAS-GFP,UAS-Wnd KD, in (E and E’) is PDF-Gal4, UAS-GFP/+; /UAS-Wnd E/ Dscam RNAi;, in (F and F’) is PDF-Gal4, UAS-GFP/+; PDF-Gal4, UAS-2x eGFP/Dscam RNAi;. Dotted insets have been zoomed in to better illustrate the diverse axonal phenotypes obtained. Scale bars are 20 μm.

**Supplemental Figure 4.**
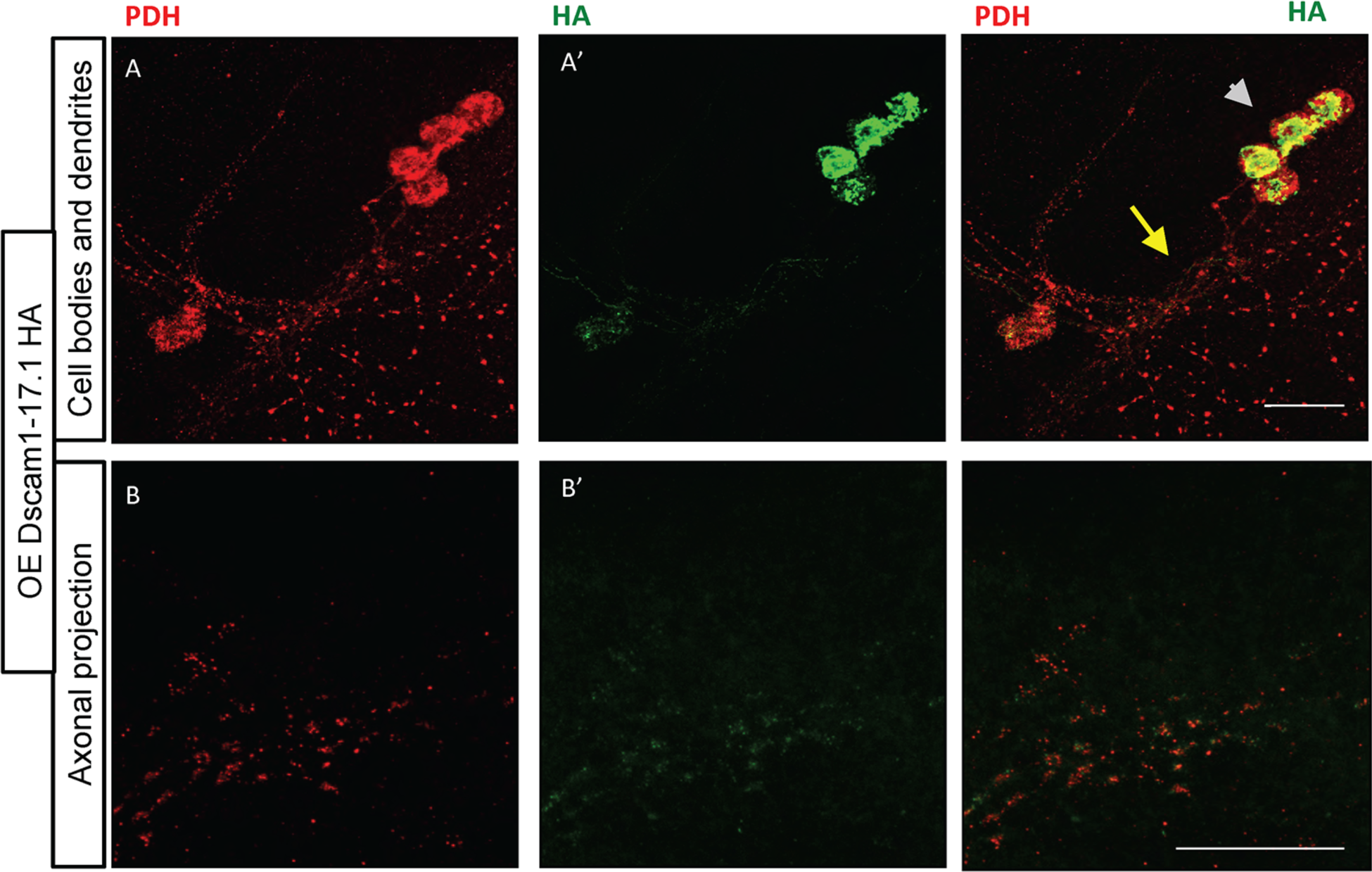
Dscam containing the TM1 domain localizes to both axonal projections as well as cell bodies and dendrites of sLNvs. **(A,B)** A Dscam form containing the TM1 domain (Dscam1-1.34.31.1 HA) localizes to both dendrites and axonal projections, and to the cell bodies. An antibody against the pigment dispersing factor hormone (PDF) specifically stains PDF neurons (A and B). Dscam expression pattern was visualized using an antibody against HA (A’ and B’). Genotype of flies is PDF-Gal4, UAS-GFP/+; PDF-Gal4, UAS-2x eGFP/+; UAS-Dscam1-1.34.31.1.HA. Scale bars are 30 μm.

---

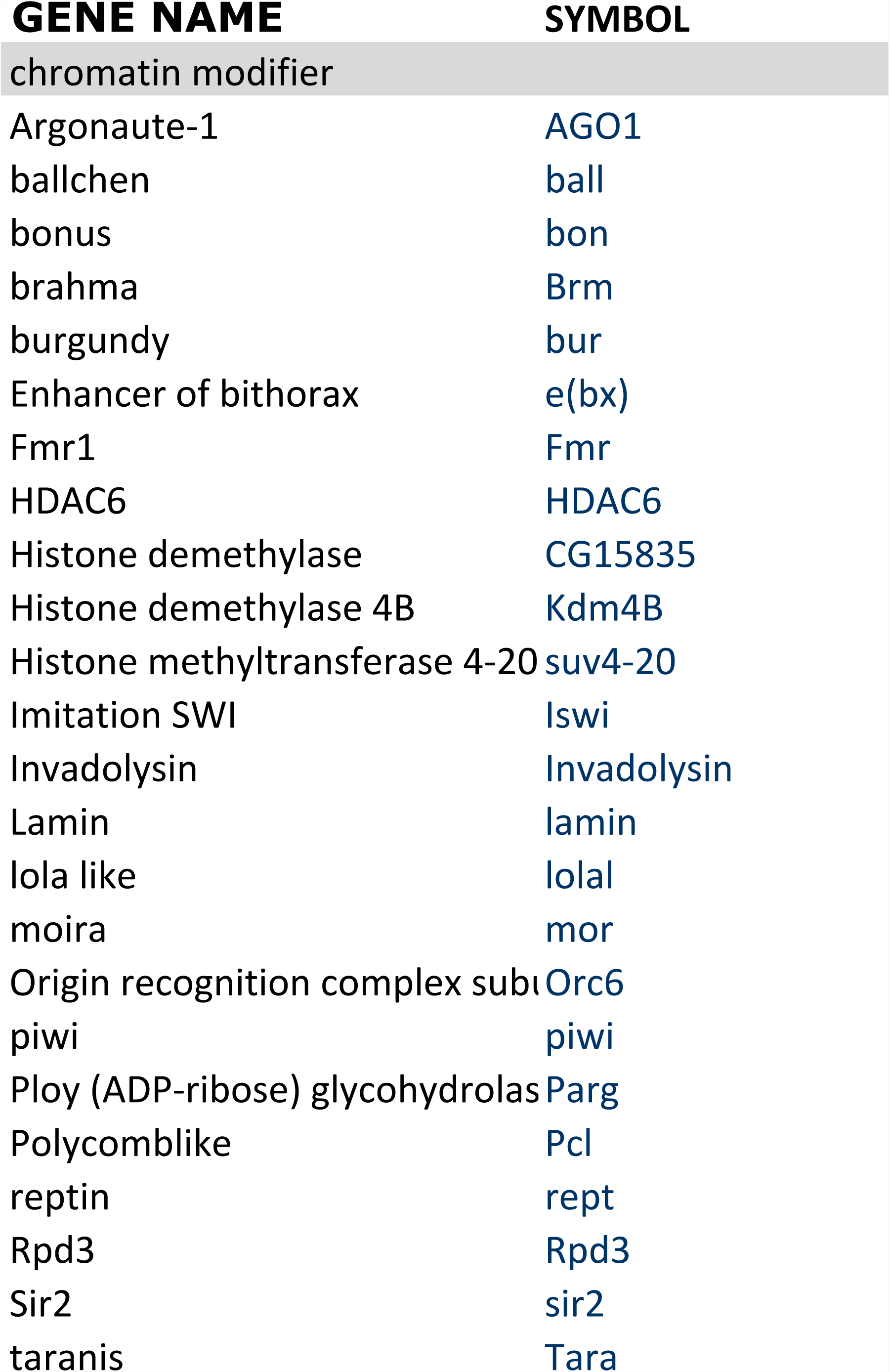

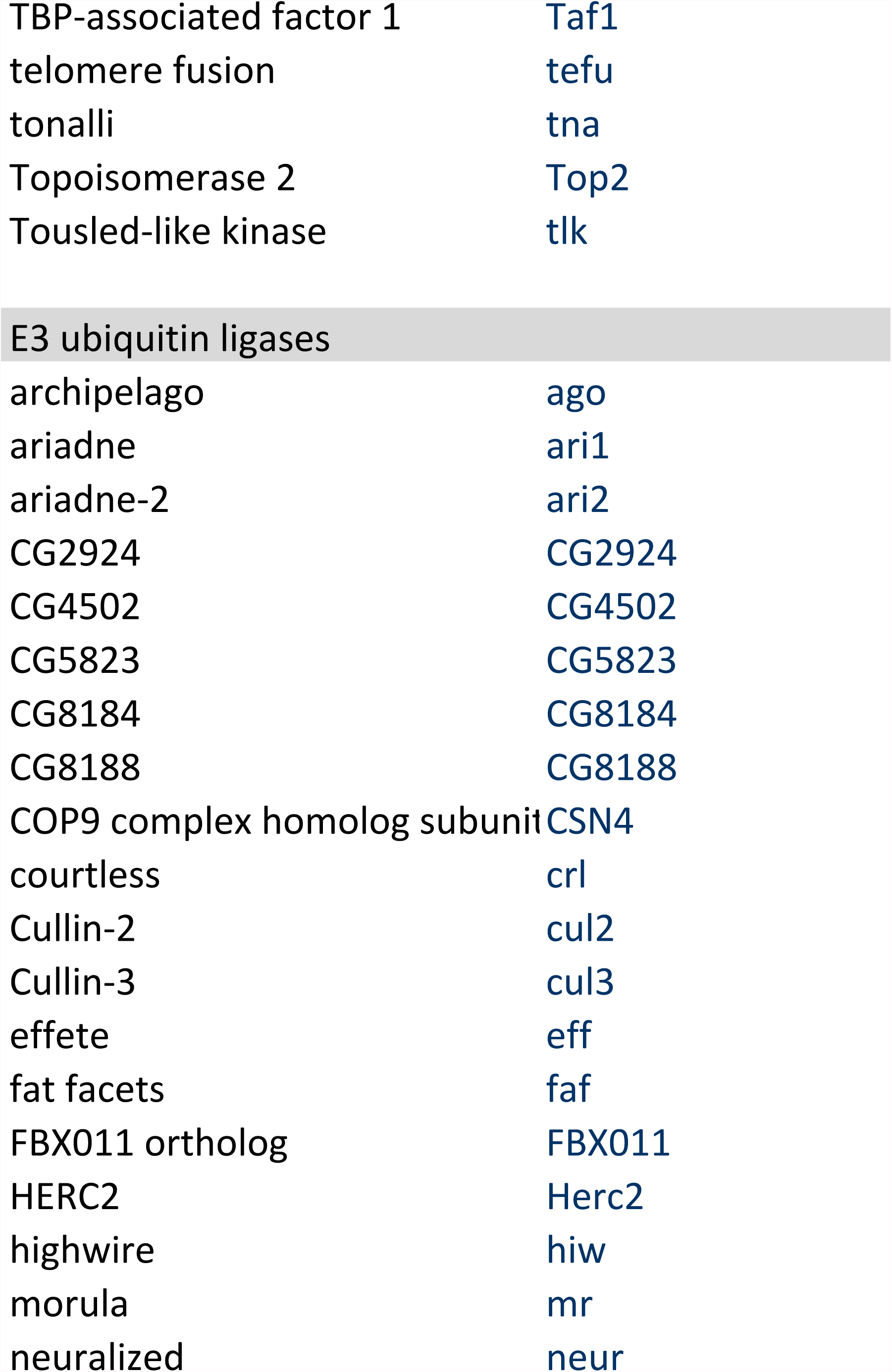

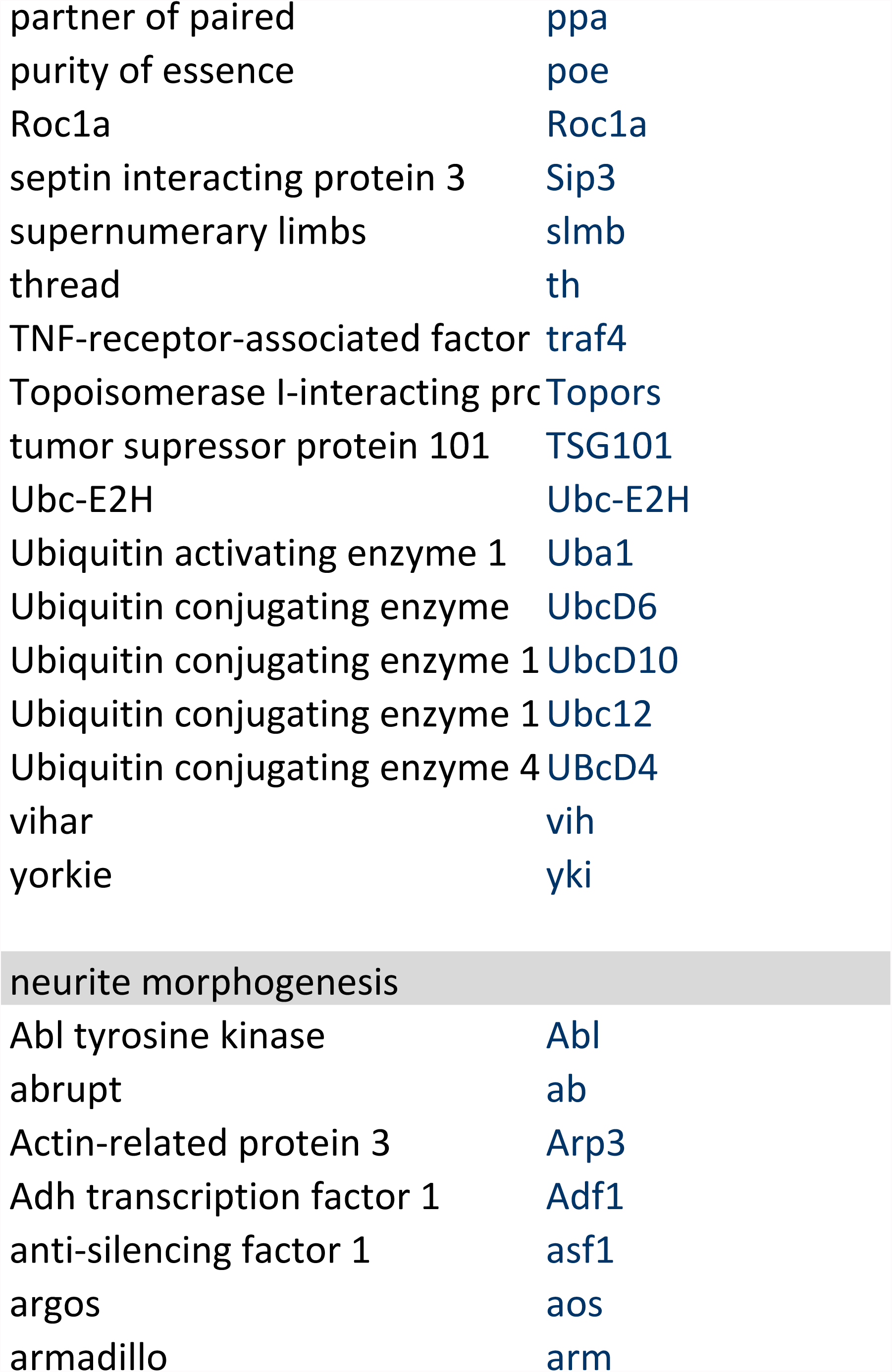

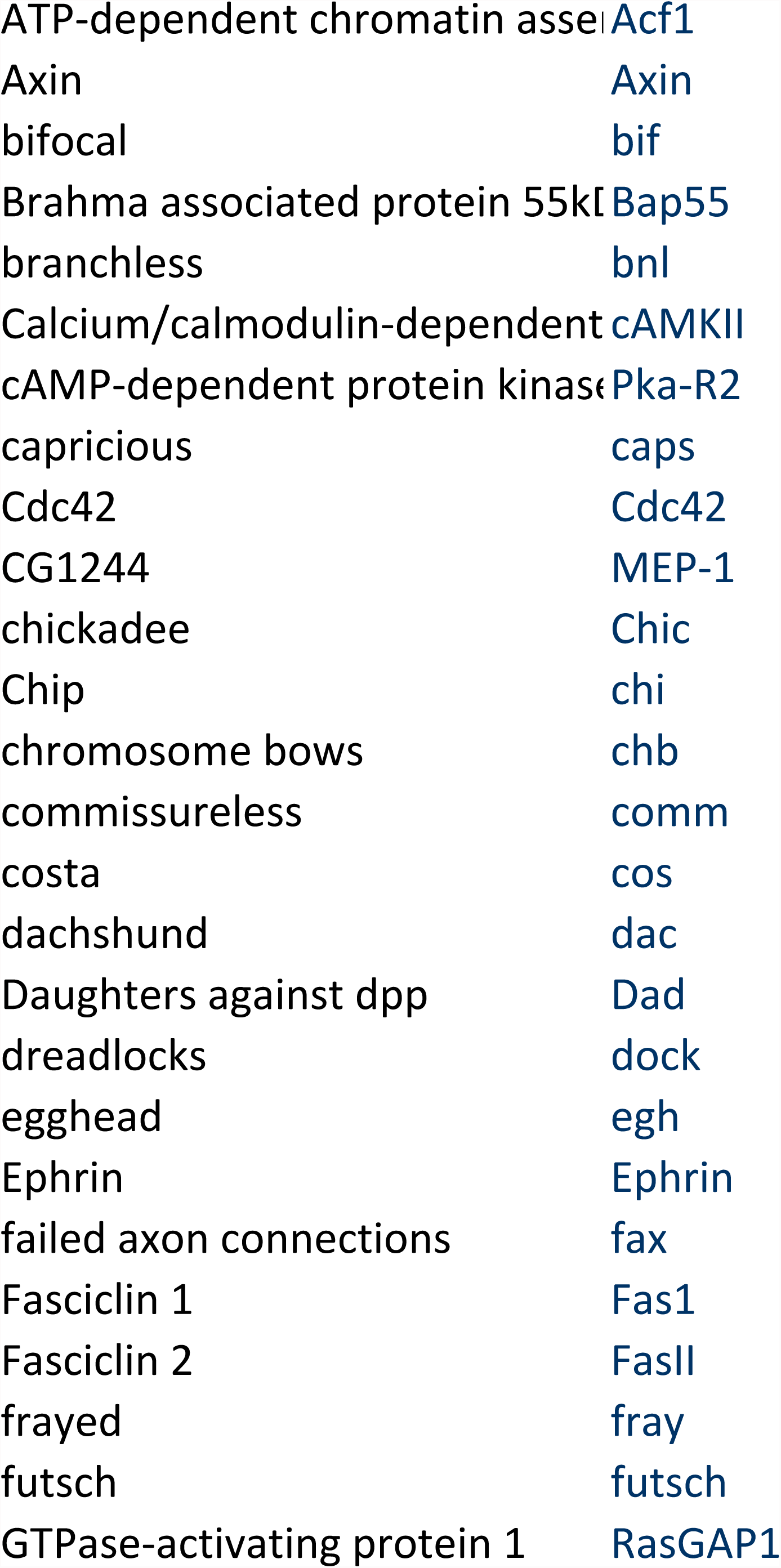

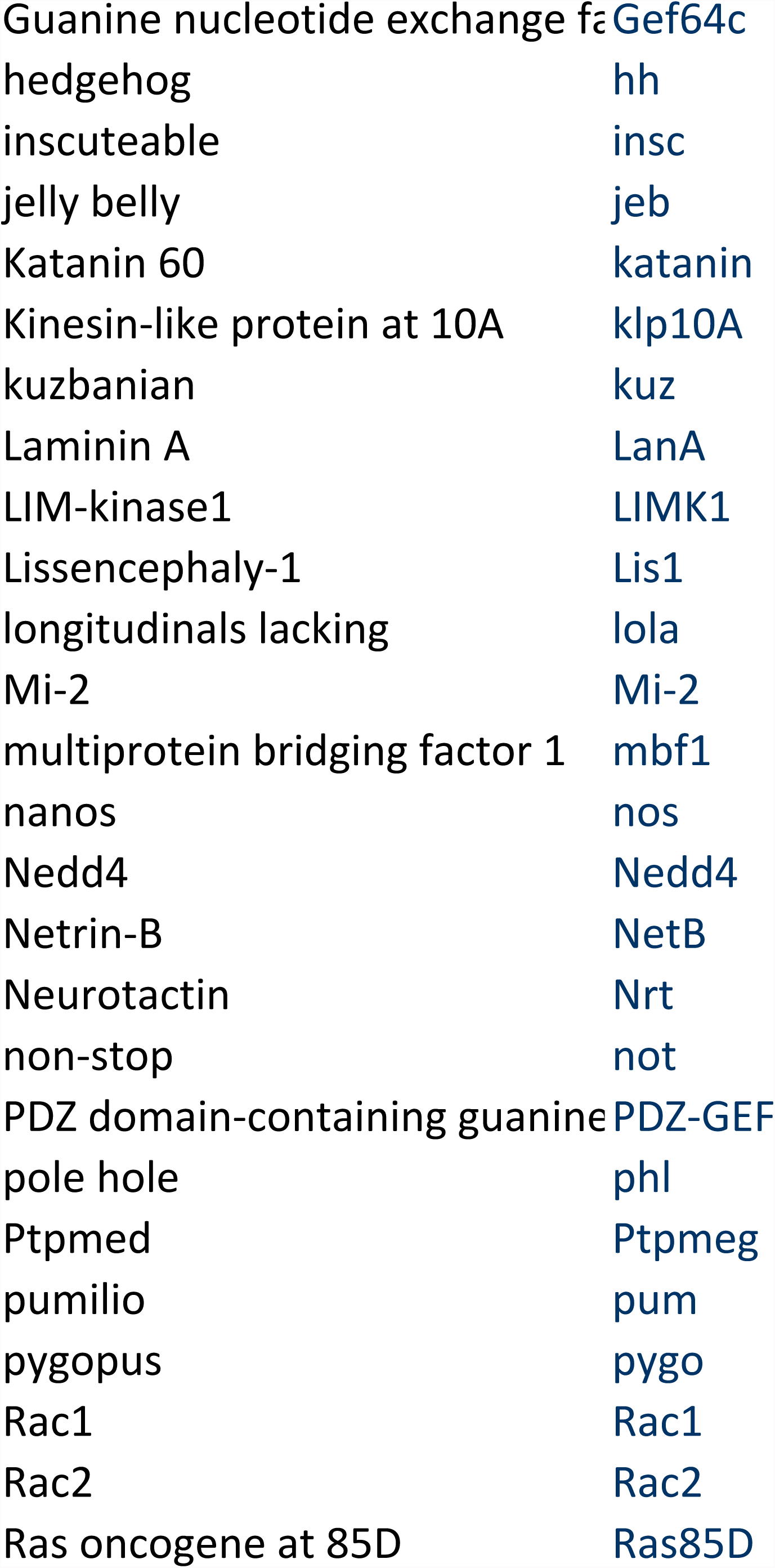

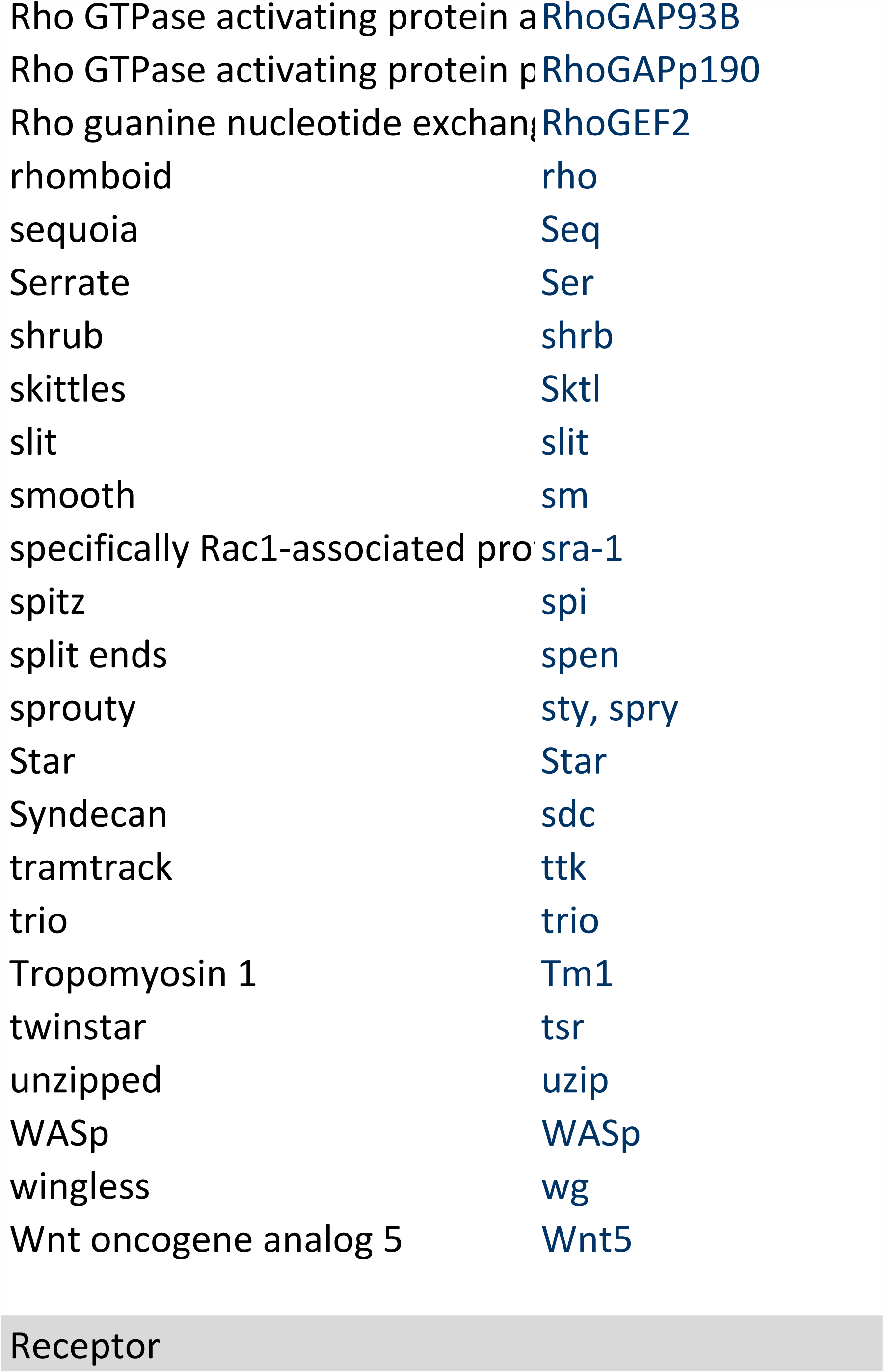

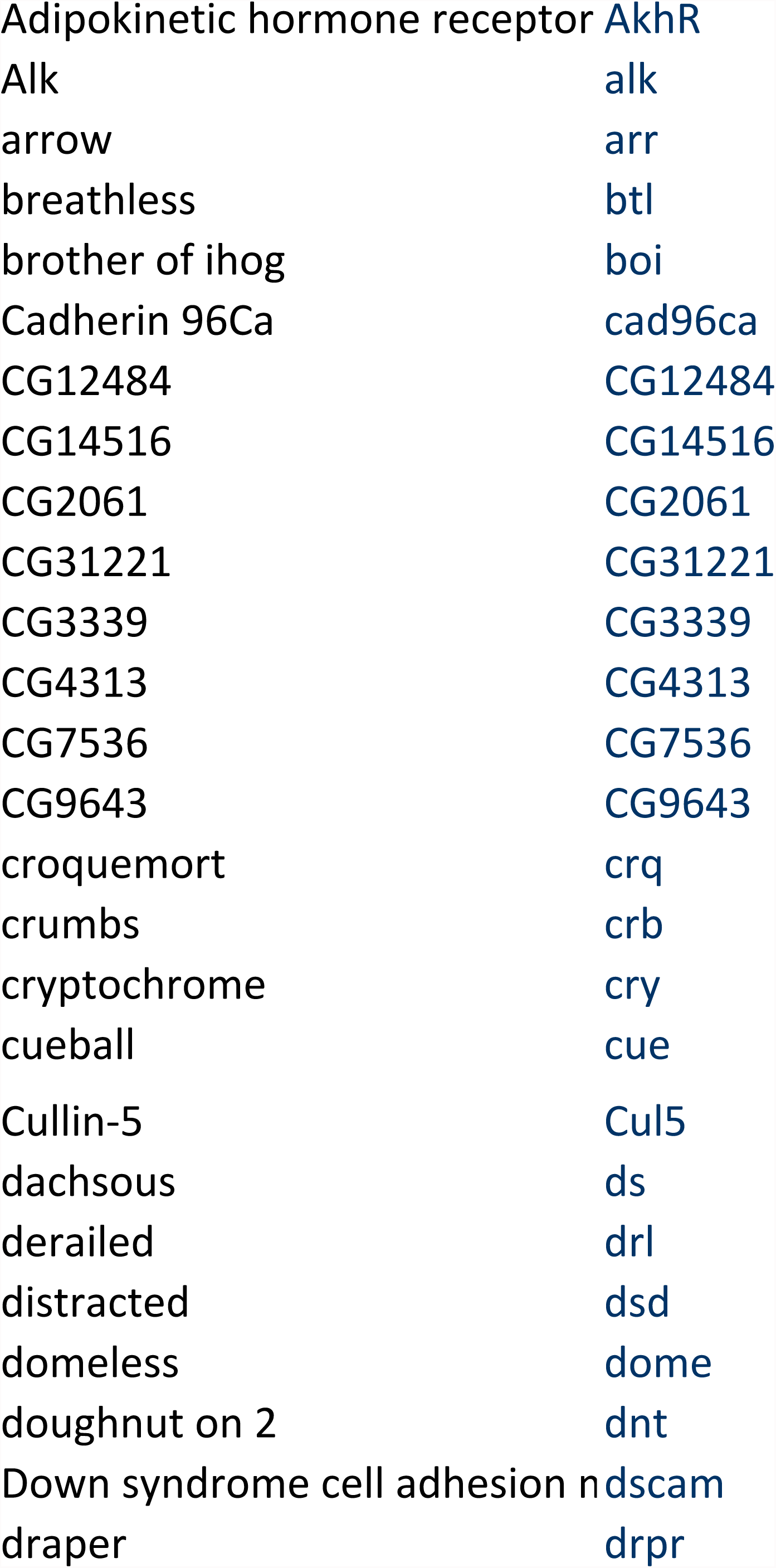

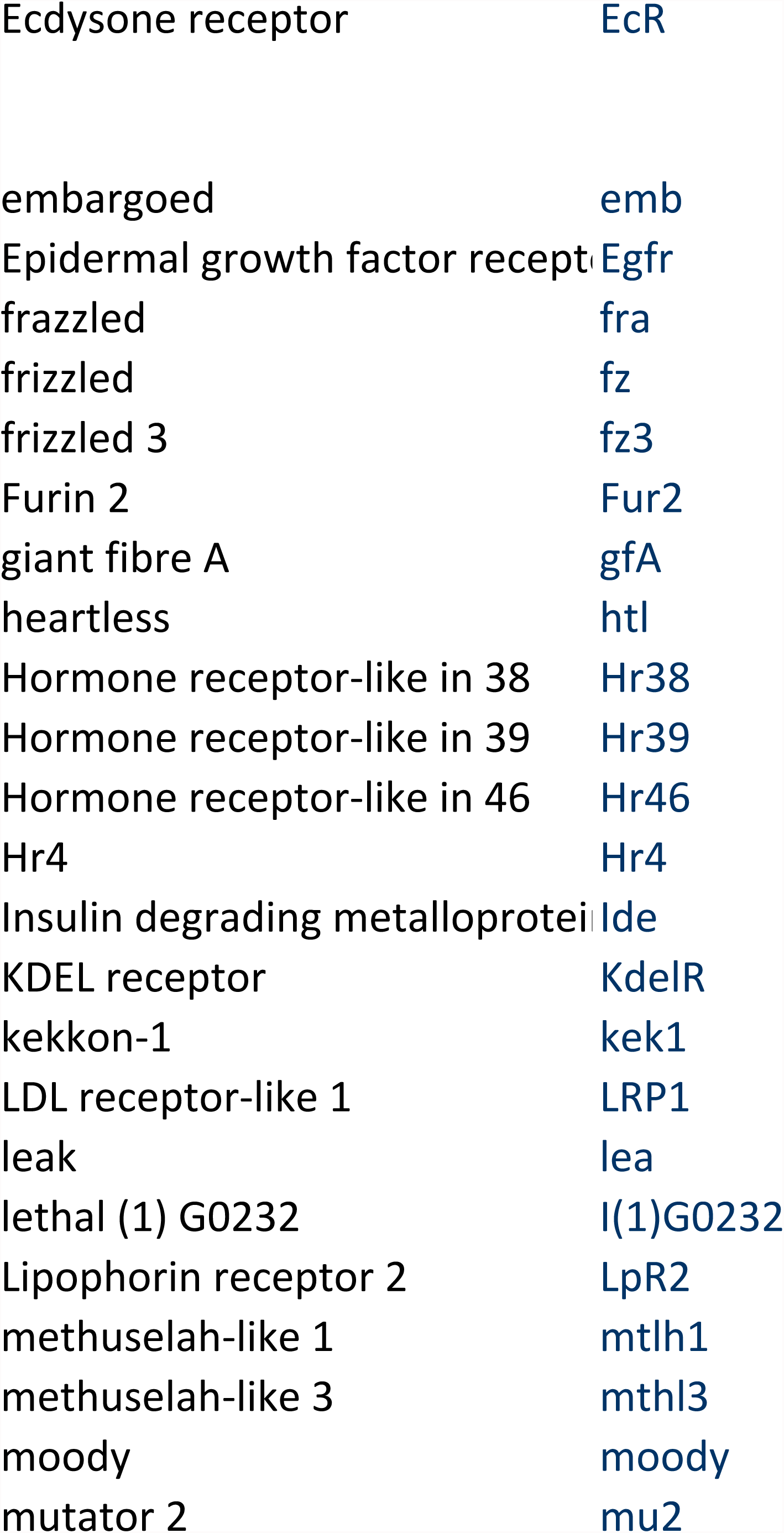

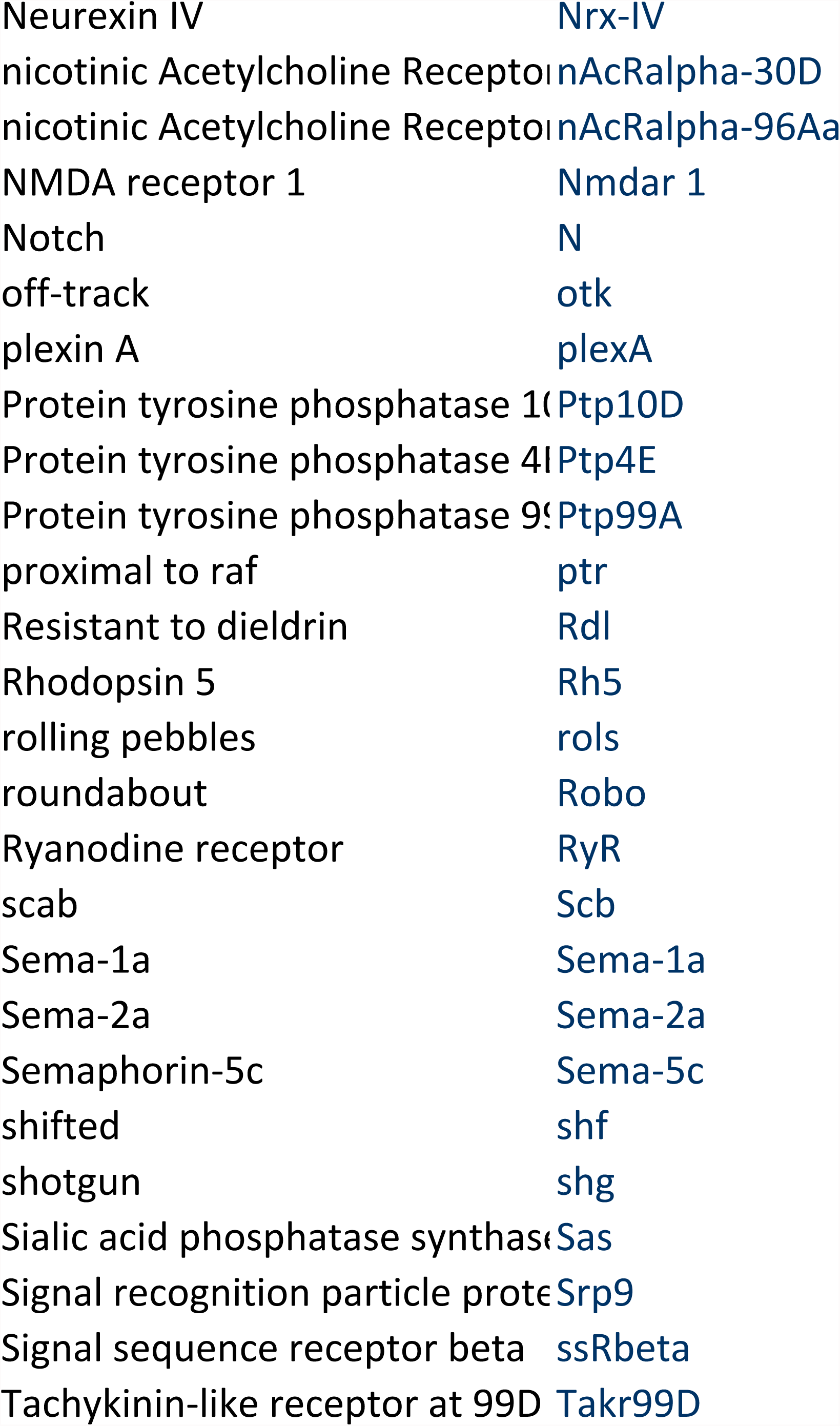

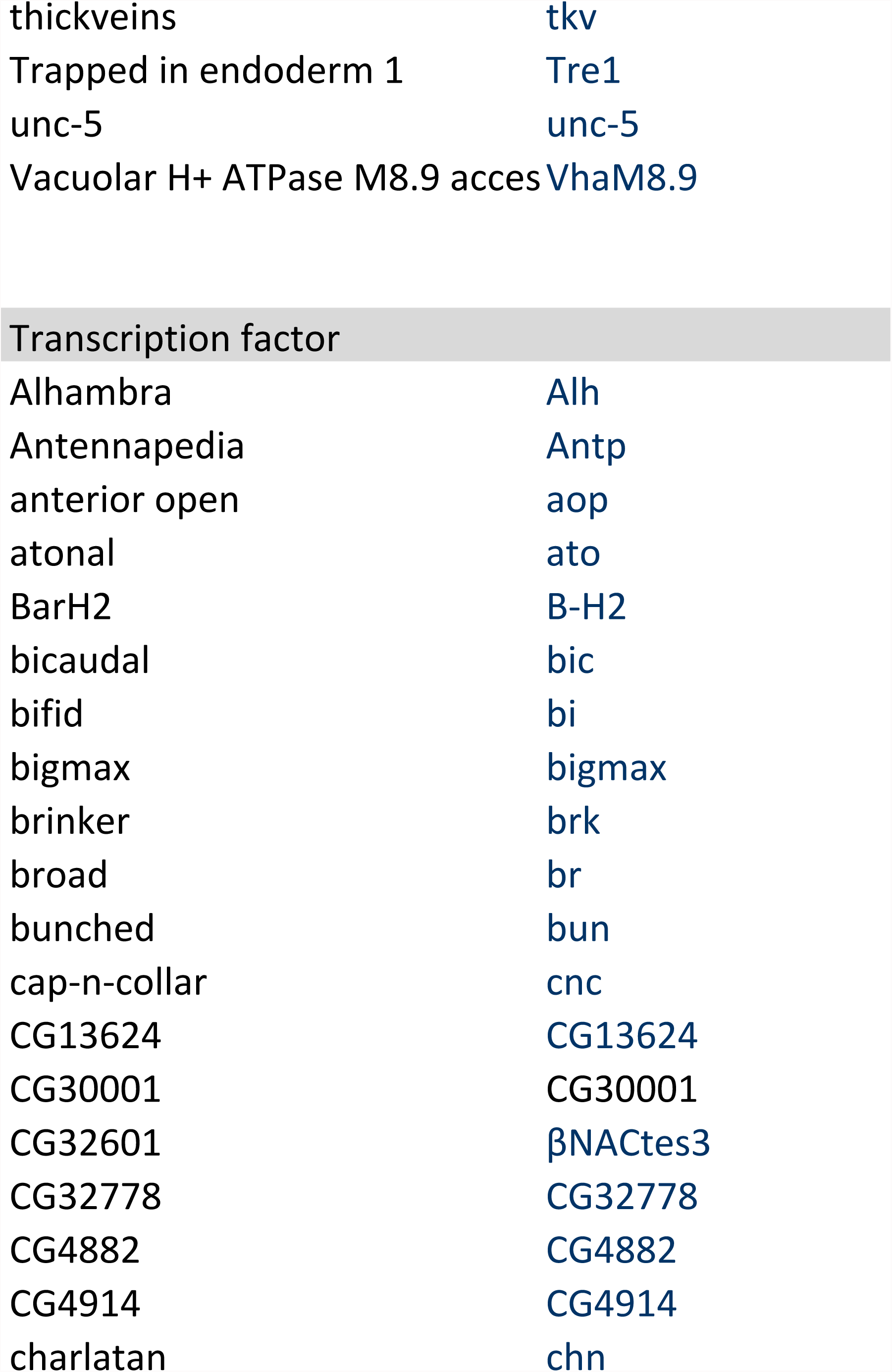

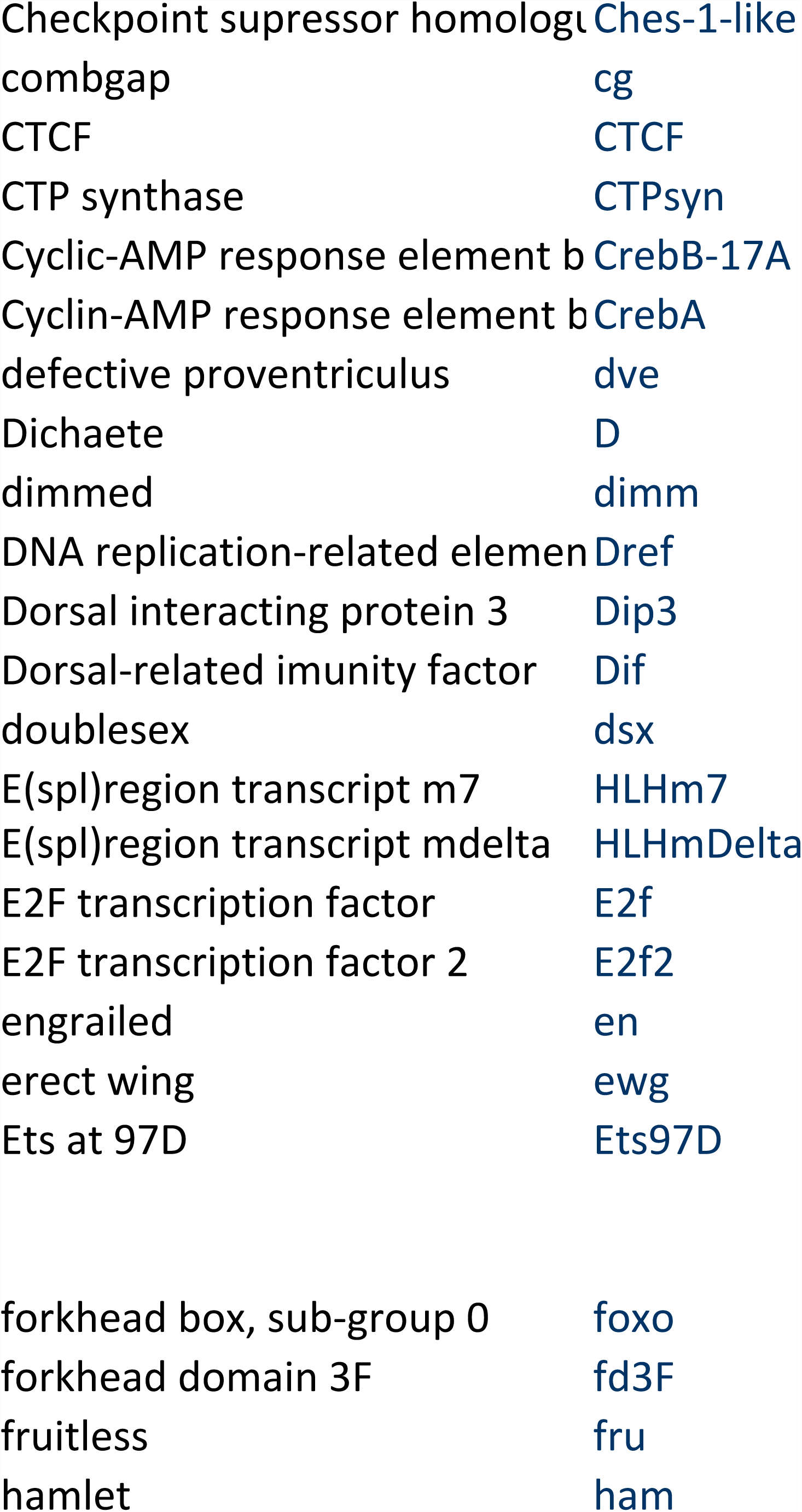

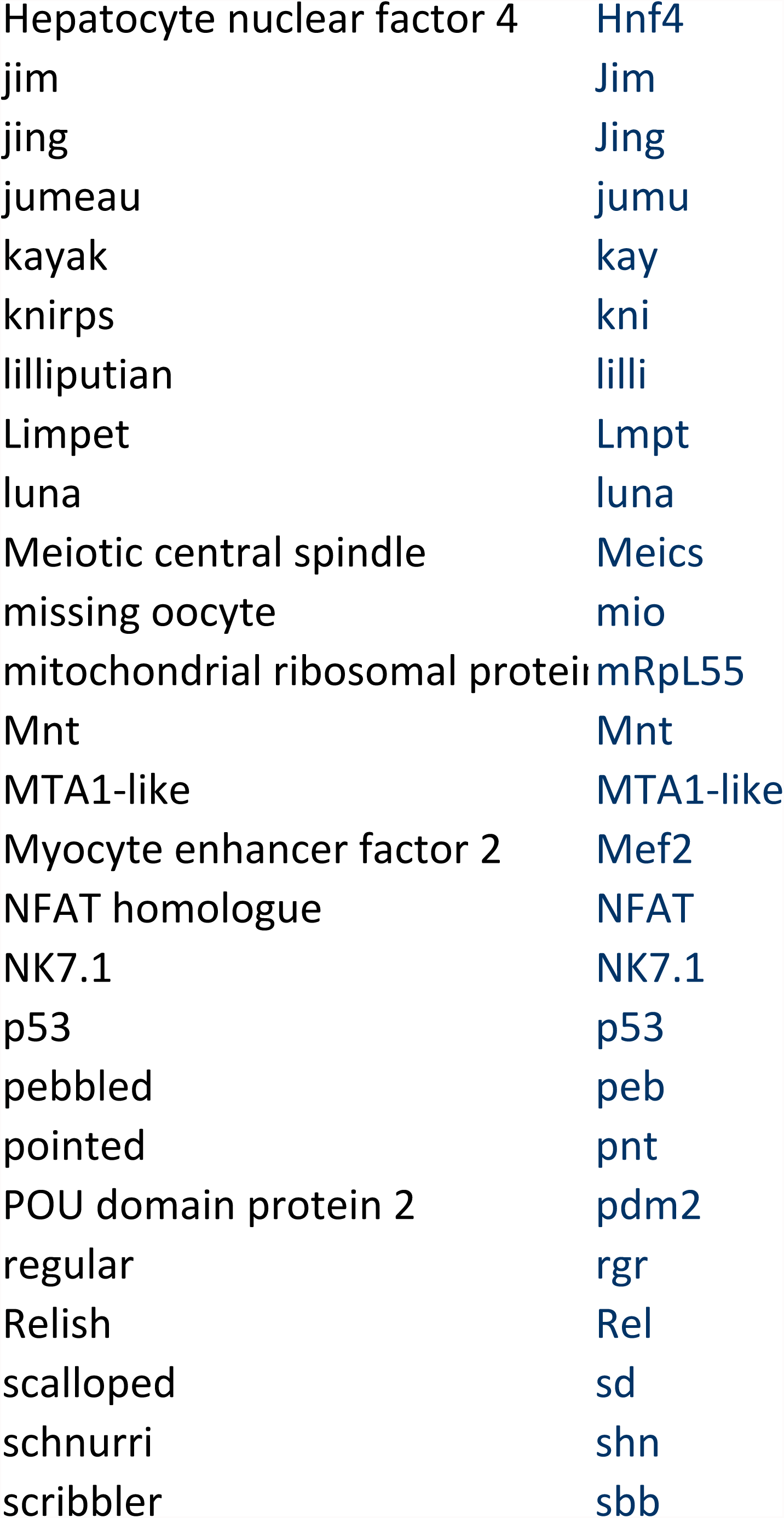

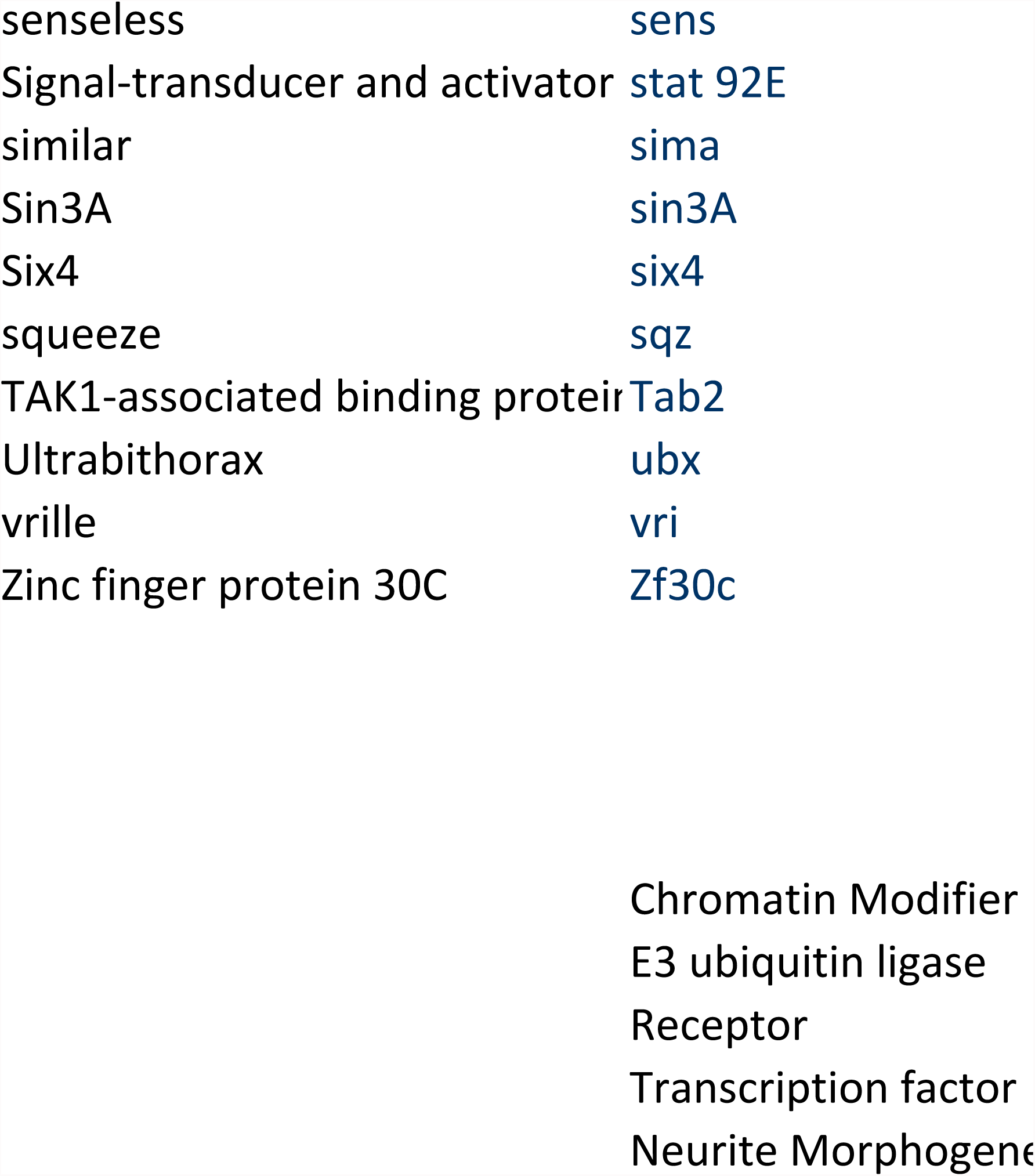

## CG number

CG6671
CG6386
CG5206
CG5942
CG9242
CG32346
CG6203
CG6170
CG15835
CG33182
CG13363
CG8625
CG3953
CG6944
CG5738
CG18740
CG1584
CG6122
CG2864
CG5109
CG9750
CG7471
CG5216
CG6889
CG17603
CG6535
CG7958
CG10223
CG34412

CG15010
CG5659
CG5709
CG2924
CG4502
CG5823
CG8184
CG8188
CG8725
CG4443
CG1512
CG42616
CG7425
CG1945
CG9461
CG11734
CG32592
CG3060
CG11988
CG9952
CG14472
CG16982
CG1937
CG3412
CG12284
CG3048
CG15104
CG9712
CG2257
CG1782
CG2013
CG5788
CG7375
CG8284
CG10682
CG4005

CG4032
CG4807
CG7558
CG15845
CG9383
CG4531
CG11579
CG1966
CG7926
CG1822
CG6546
CG4608
CG18069
CG15862
CG11282
CG12530
CG1244
CG9553
CG3924
CG32435
CG17943
CG1708
CG4952
CG5201
CG3727
CG9659
CG1862
CG4609
CG6588
CG3665
CG7693
CG34387
CG6721
CG32239
CG4637
CG11312
CG 30040
CG10229
CG1453
CG7147
CG10236
CG1848
CG8440
CG12052
CG8103
CG4143
CG5637
CG42279
CG10521
CG9704
CG4166
CG9491
CG2845
CG1228
CG9755
CG11518
CG2248
CG8556
CG9375
CG3421
CG32555
CG9635
CG1004
CG32904
CG6127
CG8055
CG9985
CG8355
CG9218
CG4931
CG10334
CG18497
CG1921
CG4385
CG10497
CG1856
CG18214
CG4898
CG4254
CG3533
CG1520
CG4889
CG6407

CG11325
CG8250
CG5912
CG32134
CG32796
CG10244
CG12484
CG14516
CG2061
CG31221
CG3339
CG4313
CG7536
CG9643
CG4280
CG6383
CG3772
CG12086
CG1401
CG17941
CG17348
CG5634
CG14226
CG17559
CG17800
CG2086
CG1765

CG13387
CG10079
CG8581
CG17697
CG16785
CG18734
CG32538
CG7223
CG1864
CG8676
CG33183
CG16902
CG5517
CG5183
CG12283
CG33087
CG5481
CG32697
CG31092
CG4521
CG6530
CG4322
CG 1960
CG6827
CG4128
CG5610
CG2902
CG3936
CG8967
CG11081
CG1817
CG6899
CG11516
CG2841
CG10537
CG5279
CG32096
CG13521
CG10844
CG809 5
CG18405
CG4700
CG5661
CG3135
CG3722
CG5232
CG8268
CG5474
CG7887
CG14026
CG3171
CG8166
CG8444

CG1070
CG1028
CG3166
CG7508
CG5488
CG3644
CG3578
CG3350
CG9653
CG11491
CG42281
CG17894
CG13624
CG30001
CG32601
CG32778
CG4882
CG4914
CG11798
CG12690
CG8367
CG8591
CG 6854
CG6103
CG7450
CG5799
CG5893
CG8667
CG5838
CG12767
CG6794
CG11094
CG8361
CG8328
CG6376
CG 1071
CG9015
CG3114
CG6338

CG3143
CG12632
CG14307
CG31753
CG9310
CG11352
CG9397
CG4029
CG33956
CG4717
CG8817
CG42679
CG33473
CG8474
CG7074
CG14283
CG13316
CG2244
CG1429
CG11172
CG8524
CG33336
CG12212
CG17077
CG12287
CG8643
CG11992
CG8544
CG7734
CG5580
CG32120
CG4257
CG7951
CG8815
CG 3871
CG5557
CG7417
CG10388
CG14029
CG3998

29

36

80

79

83

307

## Results qPCR : Dscam mRNA levels upon Faf electroporation

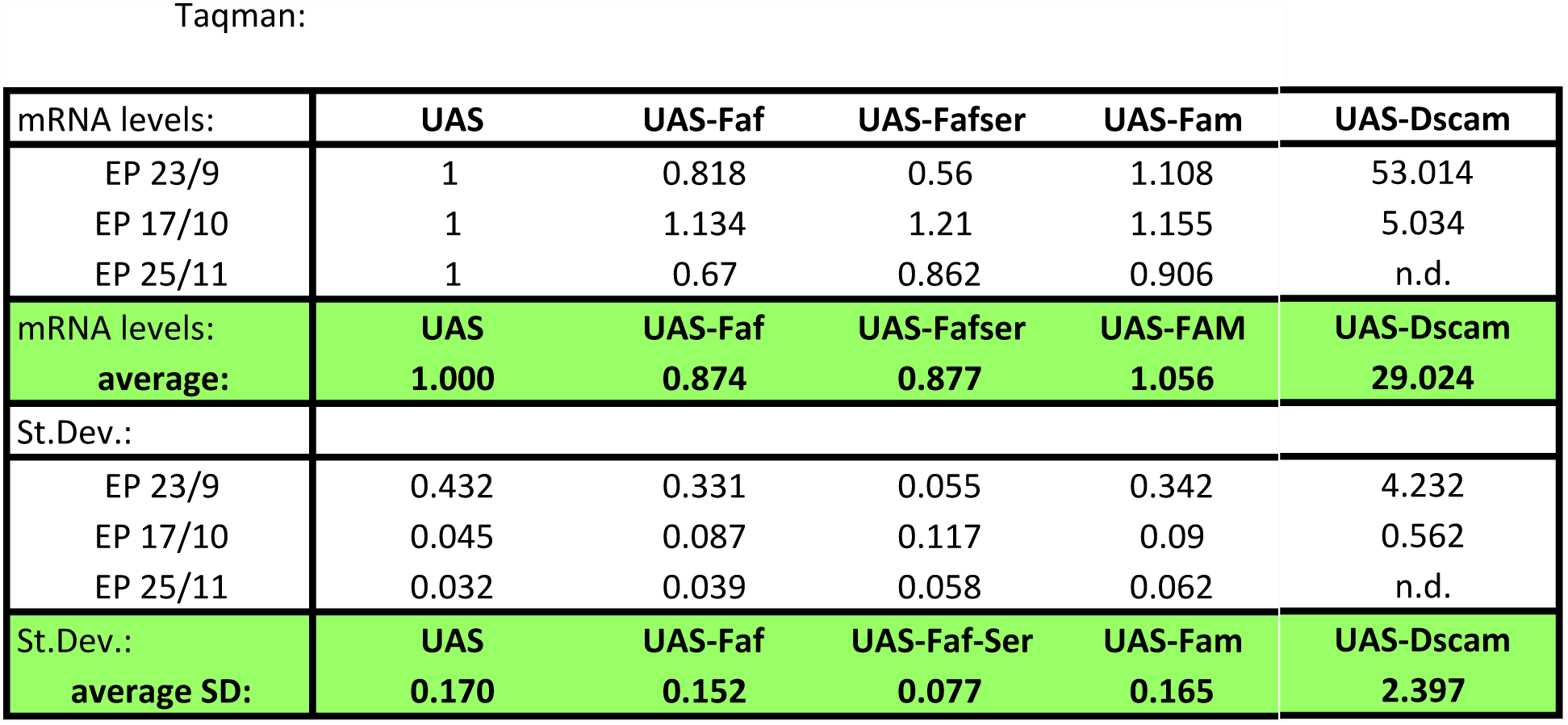

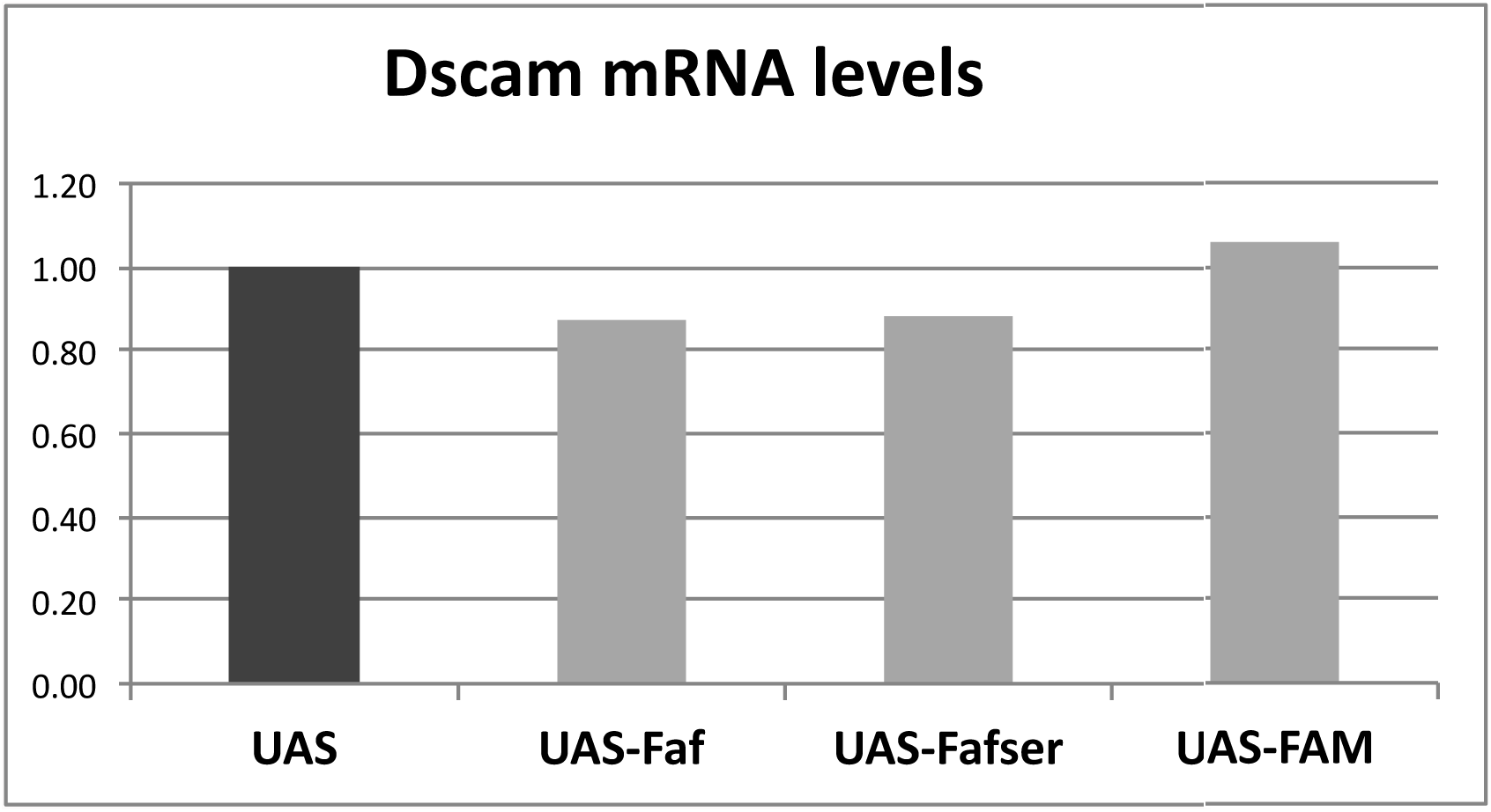

